# “Temporal control of tumor growth in nocturnal mammals: impact of the circadian system”

**DOI:** 10.1101/2020.05.25.114652

**Authors:** Paula M. Wagner, César G. Prucca, Fabiola N. Velázquez, Lucas Sosa Alderete, Beatriz L. Caputto, Mario E. Guido

## Abstract

Glioblastoma multiforme is the most aggressive brain tumor; however, little is known about the impact of the circadian system on the tumor formation, growth and treatment. We investigated day/night differences in tumor growth after injecting A530 glioma cells isolated from malignant peripheral nerve sheath tumor of NPcis (*Trp53*^*+/-*^; *Nf1*^*+/-*^) mice. Tumors generated in the sciatic nerve zone of C57BL/6 mice injected early at night in the light/dark cycle or in constant darkness, showed higher growth rates than in animals injected diurnally. Similar nocturnal increases were observed when injecting B16 melanoma cells or when mice received knocked-down clock gene *Bmal*1 cells. Moreover, treatment with a low-dose of the proteasome inhibitor Bortezomib (0.5 mg/kg) in tumor-bearing animals, displayed higher efficacy when administered at night. Results suggest the existence of a precise temporal control of tumor growth and of drug efficacy in which the host state and susceptibility are critical.

## Introduction

Carcinogenesis is a complex neoplastic process resulting in the accumulation of genetic alterations primarily in genes involved in the regulation of signaling pathways relevant to the control of cell growth and division (reviewed in [1]).

Typical characteristics of abnormal growth and proliferation include sustained proliferative activation, growth suppressor evasion, cell death resistance, replicative immortality, induction of angiogenesis, invasiveness and metastasis, all of which are based on genome instability and inflammation. A hallmark of emerging importance is avoidance of immune destruction and deregulation of cellular metabolisms. Beyond the multi-etiological causes of carcinogenesis, the advent of modern lifestyles involving longer artificial illumination exposure, hypercaloric diets, nocturnal shift work, sedentary life and jet lag, among others, severely disturbs the normal physiology and behavior of human beings. Vertebrates including mammals and particularly humans have become adapted through evolution to the natural light/dark (LD) cycle of day and night and have developed an internal clock that temporally rules physiology and behavior with a near-24 h-period (circadian). In this connection, the World Health Organization’s International Agency for Research on Cancer has classified shift work involving circadian disruption into Group 2A (probably carcinogenic to humans) [2]. Disruption of these time-regulated processes and misalignment of the internal circadian sleep-wake cycle caused by work at night, together with longer artificial illumination exposure, may predispose individuals to the development of cancer [3,4].

At the molecular level, the circadian clock operates through a transcriptional/translational feedback loop that involves a group of regulatory transcription factors -clock genes and proteins-including *Clock, Bmal1, NPAS2, Periods (Per)* and *Cryptochromes (Cry)*, among others [5,6]. Once translated, these proteins display activity as activators (CLOCK, BMAL1, NPAS2) or repressors (PER, CRY) of gene transcription and are capable of driving clock gene rhythms and rhythmic clock-controlled gene expression under a 24 h cycle through a consensus sequence E-box at the promoters of target genes [5]. A secondary cycle involves the nuclear receptor REV-ERBα/β (REV): when expression levels of this receptor are high, it binds to RORE sites present in promoter and enhancer regions of the *Clock* and *Bmal*1 genes, repressing its transcription. In contrast, when the REV-ERBs levels are low, another nuclear receptor of retinoic acid, ROR, binds to the RORE site of promoters and may activate *Clock* and *Bmal1* transcription [7,8]. *Per*1 and 2 are involved in the synchronization of the molecular clock after stimulation by early response transcription factors whose activity is controlled by extracellular signals such as hormones, second messengers, temperature or neurotransmitters. Among these transcription factors are cAMP response elements and protein-CRE binding (CREB) elements, HSF1 that binds to heat shock response elements (HSEs), serum response factors (SRF) that bind to response elements serum (SREs) and glucocorticoid receptors (GR) that bind to glucocorticoid response elements (GREs) [9]. At the cellular level, a redox / metabolic clock has been reported to drive peroxiredoxin (PRX) oxidation cycles and shown to be highly conserved through evolution. These redox oscillators are present in all kingdoms of life and even when they interact with the transcriptional molecular clock they can still function in the absence of transcription, as occurs in enucleated cells [10].

Through their rhythmic transcription, circadian clock genes regulate the metabolism and respiration of cells, therefore becoming very important regulators of the cell division cycle [3]. Loss of clock gene function or misalignment of circadian rhythms may, therefore, lead to diverse metabolic disorders (obesity, diabetes, hyperlipidemia, etc.) [11] and higher risk of cancer by inducing malignant cell growth and tumor development [3]. Animal models carrying mutations in the clock proteins exhibit premature aging, metabolic disorders, immune deficiency and proclivity for developing cancer with several cellular functions severely affected such as apoptosis, cell cycle control, chromatin remodeling, DNA damage repair, among others (reviewed in Fu & Kettner, 2013). As circadian clock genes and their clock-controlled genes are implied in a diversity of cellular functions and cancer-related biological pathways, the disruption of their gene expression may promote cancer development (reviewed in [3]). Moreover, individuals carrying certain clock gene variants can be more susceptible to the development of cancer when exposed to prolonged circadian disruption and light exposure at night during sustained shift work activities.

In this context, a deeper understanding is required of how circadian biology can significantly contribute to the knowledge and potential treatment of human pathologies, in particular cancer and metabolic and behavioral disorders [3,13].

Brain tumors are very aggressive, particularly those of glial origin. Gliomas include astrocytomas, oligodendrogliomas, and glioblastomas and vary in their aggressiveness or malignancy. Some are slow-growing and are likely curable. Others are fast-growing, invasive, difficult to treat, and likely to recur [14]. In this respect, the glioblastoma multiforme (GBM) is particularly aggressive and with a very poor prognosis. Attempts have been made to replicate this neoplastic pathology of humans in animal models [15,16]. One such model is generated by inoculating A530 glioma cells isolated from the malignant peripheral nerve sheath tumor (MPNSTs) of NPcis (*Trp53*^*+/-*^; *Nf1*^*+/-*^) mice, a model of human neurofibromatosis type I [17]. A530 cells were firstly characterized in culture and then injected in peripheral nerves of C57BL/6 mice to generate tumors *in vivo*. Here, we show that tumors generated by the injection of malignant cells (gliomas A530 or melanoma B16 cells) exhibit a higher rate of growth when the inoculation occurs at night. We also investigated the role of the molecular circadian clock in tumor formation and the time-regulated susceptibility to chemotherapy treatment with the proteasome inhibitor Bortezomib (BOR) [18]

## Results

### Isolation of glioma-derived A530 cells from a malignant peripheral nerve sheath tumor (MPNST)

In order to study day/night differences in the tumor growth rate after the injection of cancer cells into C57BL/6 mice, tumor cells from malignant peripheral nerve sheath tumor (MPNSTs) developed in NPcis (*Trp53*^*+/-*^; *Nf1*^*+/-*^) mice in a C57BL/6 genetic background were isolated and characterized as described in Materials and Methods (Suppl. Figs. 1-4).

### *In vivo* experiments: injection of A530 cells at the beginning of the day or night

To investigate day/night differences in the tumor growth rate after the injection of A530 cells, we carried out a series of *in vivo* experiments. For this, A530 cells characterized as described under Materials and Methods (Suppl. Figs. 1-4), were injected into the sciatic nerve zone of C57BL/6 mice at the beginning of the day (ZTs 1-2) or night (ZTs 12-13) in animals subjected to a regular 12-h light/dark (LD) cycle. Tumors were palpable by day 15 post-injection and the growth rate was measured periodically until days 28-30 post-injection (Suppl. Figure 5). Tumors growing in animals that were injected at the beginning of the night showed a higher growth rate than those in animals injected at the beginning of the day (Fig. 1A). The statistical analysis revealed a significant effect of time (p < 0.0003 by t-test with Welch-s correction). Similar results were observed when animals were synchronized by a regular LD cycle and released to constant darkness (DD) for 72 h prior to the injection of tumor cells. A higher growth rate was observed in mice injected at the beginning of the subjective night (ZTs 12-13) compared with those injected at the beginning of the subjective day (ZTs 1-2) (p < 0.047 by Mann-Whitney) (Fig. 1B). Under both illumination conditions tested (LD or DD), the doubling time index (DT) reflecting the time needed for a tumor to double in size and calculated as described in [19], was significantly lower at night (p < 0.015 by t-test) (Fig. 1B, D). Furthermore, the relative tumor volume was greater during the last days post-injection for those animals inoculated at night (Suppl. Fig. 6A-B). These results strongly suggest the existence of endogenous circadian control of tumor growth. By contrast, no differences were observed when animals were injected at the same time but with tumor cells coming from cultures synchronized at different times, 12 h apart with respect to the culture medium exchange (Fig. 1E-F) (N.S, p = 0.27). No differences were found in the relative volume of the tumor between the two groups with different cell synchronization times as shown in Suppl. Fig. 6C. These results strongly indicate that the day/night phenomenon is mainly dependent on the host state.

**Fig. 1:**
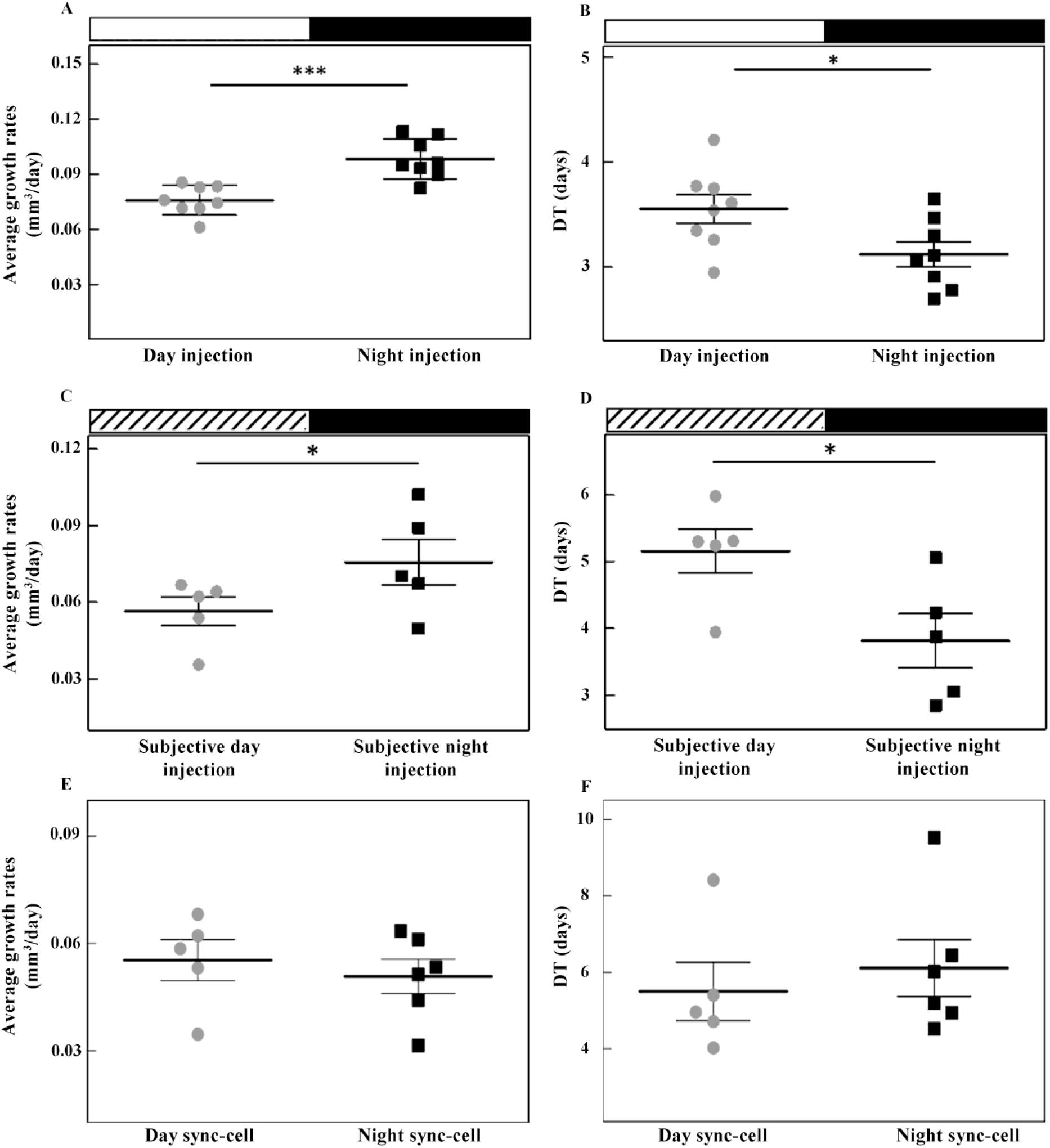
Tumor growth rate after A530 cell injection in C57BL/6 mice at the beginning of day or night. **A-B** Synchronized A530 cells (1 × 10^5^ cells in 10 µl of PBS) were injected at the beginning of the day (gray circle, n = 8) or night (black square, n = 8) into the sciatic nerve zone of C57BL/6 mice subjected to a regular LD cycle. Results revealed a significant difference in tumor growth with a higher growth rate (A, left panel) and lower values of Doubling Time (DT) index (B, right panel) in tumors growing in animals injected at the beginning of the night compared with those subjected to diurnal injection (***p < 0,0003 and *p < 0.015 by t-test with Welch’s correction, respectively). Results are mean ± SEM of two independent experiments. **C-D**. Synchronized A530 cells (1 × 10^5^ cells in 10 µl of PBS) were injected in the sciatic nerve zone of C57BL/6 mice maintained in DD at the beginning of the subjective day (gray circle, n = 5) or night (black square, n = 5). Results showed a higher growth rate (C, left panel) and lower values of DT (D, right panel) when animals were injected at the beginning of the subjective night compared with those injected at the beginning of the subjective day (*p < 0.047 and *p < 0.016 by Mann-Whitney, respectively). Results are mean ± SEM of two independent experiments. **E-F**. C57BL/6 mice were injected at the same time with A530 cells synchronized at different times with a 12 h difference after culture medium exchange (day-sync cell: gray circle, n = 5; night-sync cell: black square, n = 6). No significant differences were found in the tumor growth rate (E) or DT (F) between groups (p = 0.27 by Mann-Whitney). Results are mean ± SEM of one representative experiment. In all experiments, tumor growth rate and the DT index were calculated as described under Materials and Methods.

### *In vivo* experiments: injection of B16 cells at the beginning of the day or night

To investigate whether the finding of day/night differences in the growth rate of tumors generated by injection of glioma-derived A530 cells can be extended to other tumor models, we carried out similar *in vivo* experiments using melanoma B16 cells. These cells, previously synchronized by culture medium exchange, were injected at the beginning of the day (ZTs 1-2) or night (ZTs 12-13) into male C57BL/6 mice subjected to a regular LD cycle. We found that tumor growth exhibited higher values in animals injected at the beginning of the night (p < 0.028 by Mann-Whitney) (Fig. 2A). The DT index was significantly lower in tumor cells injected at night compared with those injected during the day (p < 0.03 by t-test) (Fig. 2B) while levels for the relative tumor volume were significantly higher 16 days post-injection in night-injected mice (p < 0.05 by two-way ANOVA) (Suppl. Fig. 6E).

**Fig. 2:**
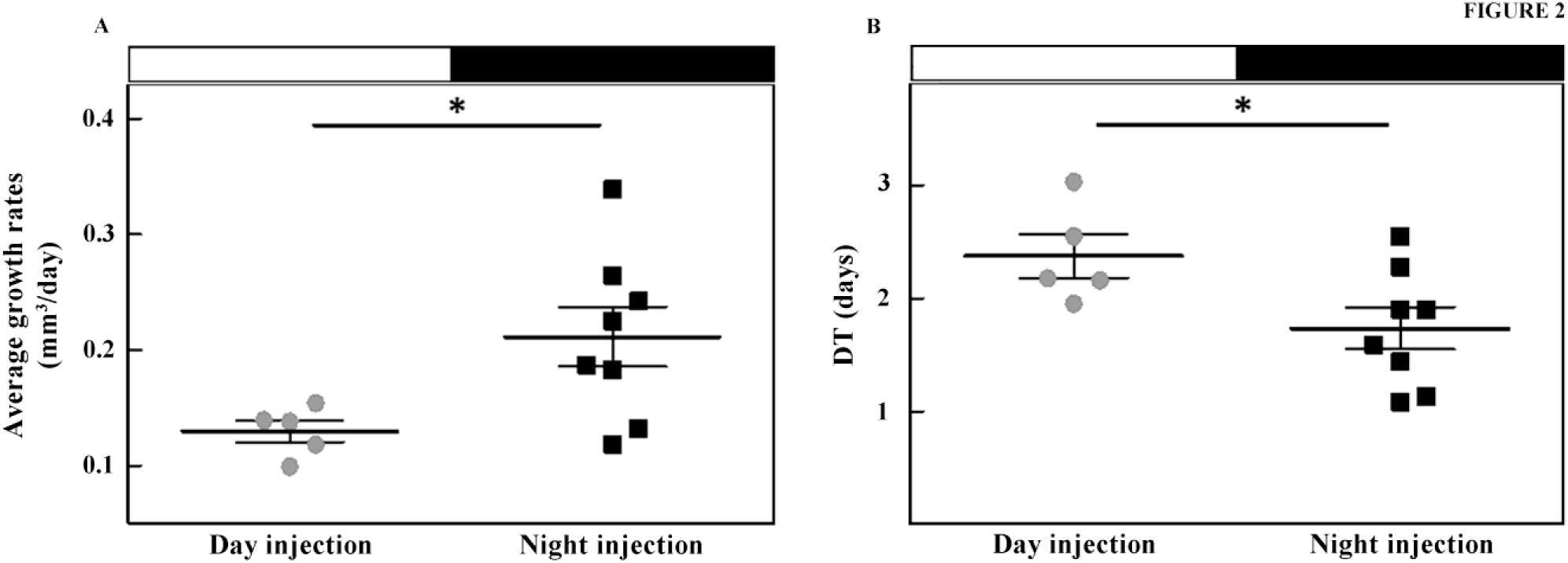
Tumor growth rate after melanoma B16 cells injection at the beginning of the day or night. Synchronized B16 cells (5 × 10^5^ cells in 50 µl of PBS) were injected subcutaneously at the beginning of the day (gray circle, n= 5) or night (black square, n = 8) in male C57BL/6 mice subjected to a regular LD cycle. Tumor growth rate and the DT index was calculated as described in Methods. Results showed a significantly higher tumor growth rate (A) and lower values of DT (B) in males of the night group compared with animals injected at the beginning of the day (*p < 0.028 by Mann-Whitney). Results are mean ± SEM of two independent experiments.

### The role of the molecular clock

To investigate the role of the molecular clock in tumor growth, we used CRISPR/Cas9 technology to knock down the expression of the molecular clock activator *Bmal*1 in A530 cells (A530 A5 cells). The disruption of *Bmal*1 gene expression was checked by RT-PCR and found to be at least 50% lower than in A530 control cells (Suppl. Fig. 7A). As a consequence, a similar effect was observed on the expression of the downstream target gene *Per*1 (Suppl. Fig. 7A). Additional effects of *Bmal*1 down regulation were found in cell cycle phases analyzed by flow cytometry (Suppl. Fig. 7 B-C). A higher proportion of A530 A5 cells in S-G_2_/M phases (59.7% for A5 and 38.7% for A530) and a higher expression of the *Bcl-2* mRNA which codifies for an antiapoptotic protein (Suppl. Fig. 7A) were observed. *In vivo* experiments showed a higher tumor growth rate in mice injected with A530 A5 cells than in animals injected with A530 control cells (Fig. 3A) (p < 0.01 by Mann-Whitney). Furthermore, the DT revealed a significant difference between the two groups (p < 0.012 by t-test): A530 control cells showed higher values than those in *Bmal1*-knocked down (KD) cells (A530 A5 cells) (Fig. 3 B).

**Fig. 3:**
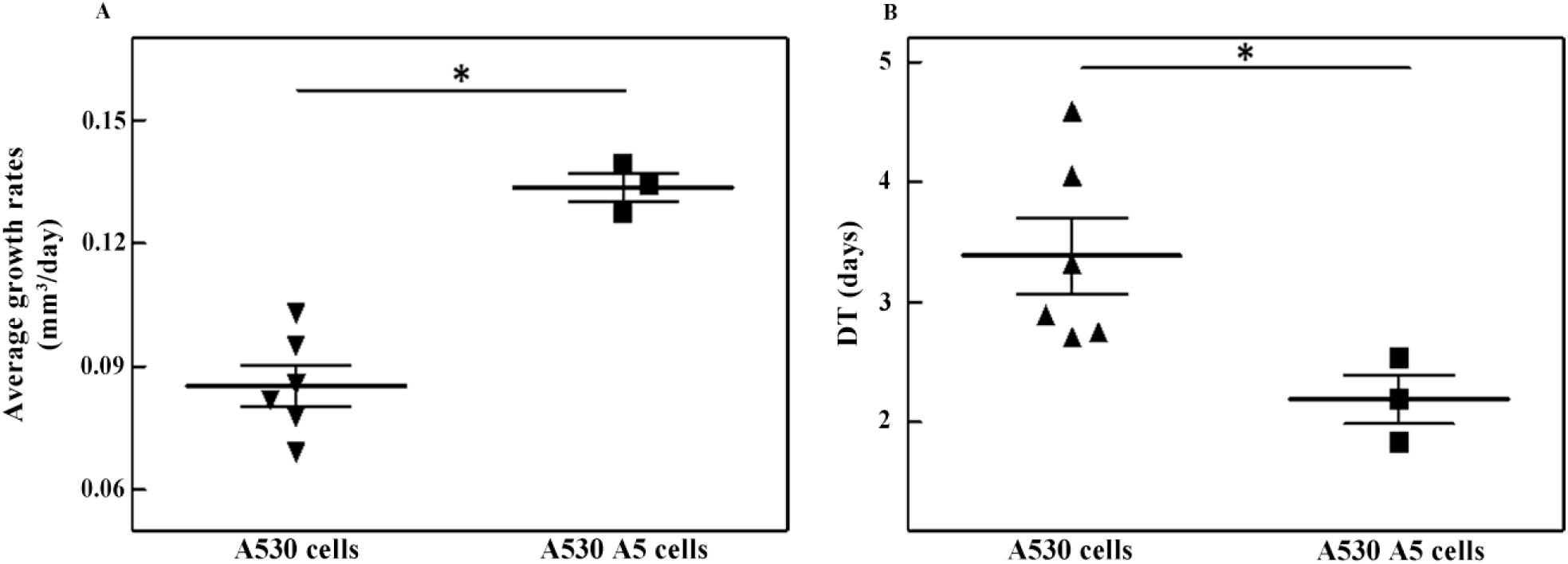
Tumor growth in animals injected with A530 or A530 A5 (*Bmal*1 knocked-down) cells. A530 cells were transfected with PX459-*Bmal1*plasmid (A530 A5 cells) to disrupt *Bmal*1 expression and injected into the sciatic nerve zone of C57BL/6 mice kept under a regular LD cycle. Results revealed a higher tumor growth rate (A) and lower values of DT (B) when mice were injected with A530 A5 cells (black square, n= 3) with decreased *Bmal*1 expression compared with A530 cells (gray circle, n = 6) (*p < 0.01 by Mann-Whitney). Tumor growth rate and the DT index were calculated as described in Methods.

### Expression of clock and clock-controlled genes in tumor and surrounding tissues

In another series of experiments, we evaluated gene expression in tumors and tissues surrounding the injection site in animals injected with A530 cells during the night (dark phase) by qPCR as shown in Fig. 4. We assessed expression of clock genes *Bmal*1 and *Per*1, the clock-controlled gene *Chokα*, and *Pcyt*2, the gene encoding for a phosphatidylethanolamine-synthesizing enzyme. Results showed no significant differences in *Bmal*1 mRNA levels between tumor and surrounding tissues in animals injected at night; however, levels of *Per*1 mRNA were substantially reduced in tumors compared with adjacent tissues (p < 0.009 by Mann Whitney) (Fig. 4A-B). By contrast, a significant increase in mRNA levels of *Chokα* was found in tumors compared with levels in the tissues surrounding the injection site (p < 0.0023 by Mann Whitney) (Fig. 4C). In addition, no significant differences were observed between mRNA levels of *Pcyt*2 in the two types of tissue (tumor and adjacent tissues) (Fig. 4D).

**Fig. 4:**
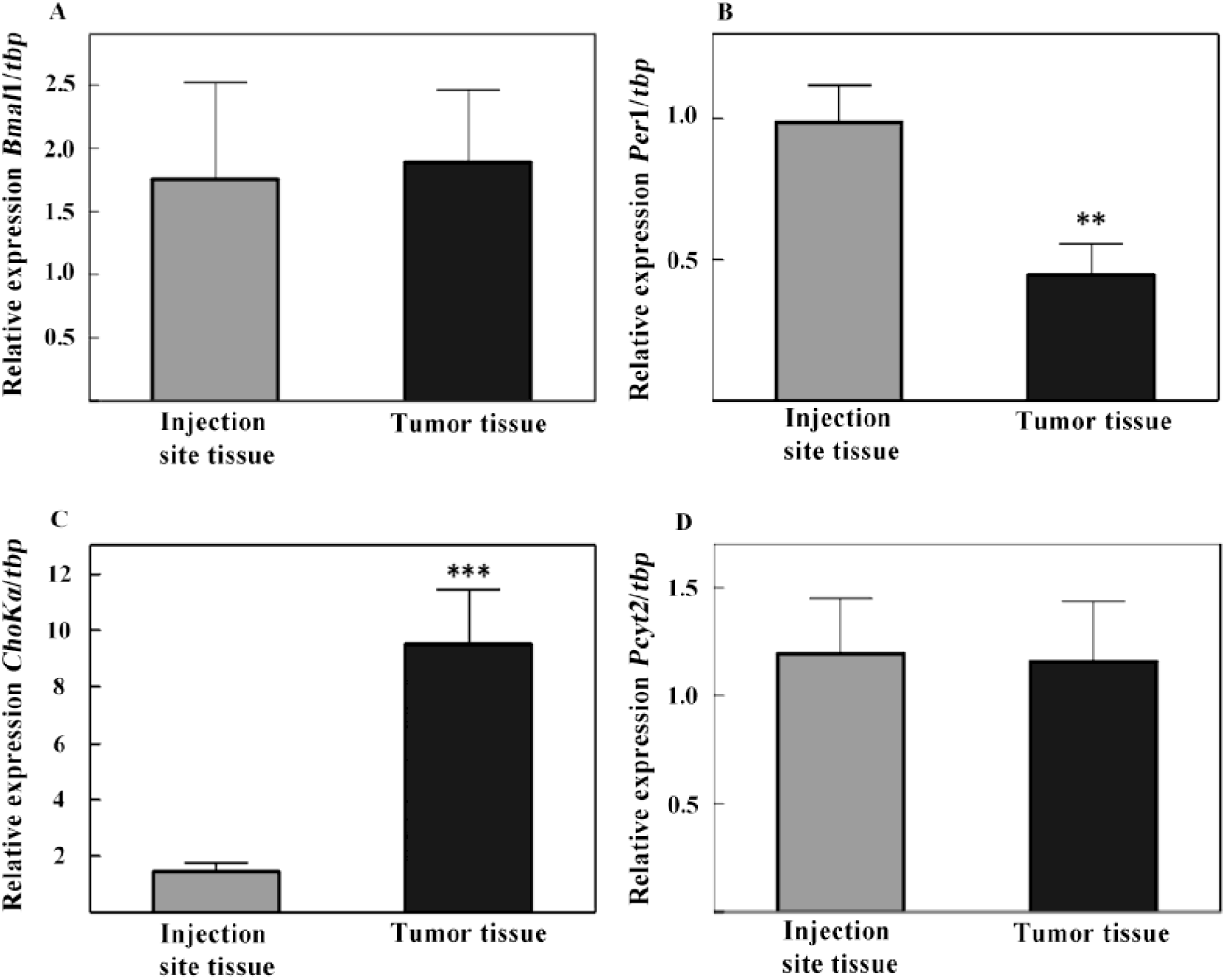
Expression of clock genes *Bmal*1 (A) and *Per*1 (B), clock-controlled gene *Chok*α (C) and lipid synthesizing enzyme gene *Pcyt*2 (D) in tumor tissue and adjacent tissue at the injection site collected from animals injected with A530 cells at the beginning of the night. mRNA levels were assessed by RT-qPCR with RNA extracted from tumor (black histograms) or adjacent tissues (gray histograms) collected from mice injected with A530 cells at the beginning of the night and sacrificed at days 28-30 post-injection. Values were normalized according to the expression of the housekeeping gene *tbp*. Results showed a significant decrease in levels of *Per*1 mRNA expression (B) in tumor tissue (black) compared with injection site tissue (gray) (**p < 0.009 by Mann-Whitney) and a significant increase in those for *Chokα* mRNA (C) expression in tumor tissue compared with control injection site tissue (**p < 0.0023 by Mann-Whitney). No differences were found in *Bmal*1 (A) and *Pcyt*2 (D) mRNA levels between both tissues. See text for further detail.

### Temporal changes in susceptibility to chemotherapy treatment with Bortezomib

The differential temporal susceptibility to chemotherapy in cell culture of A530 cells has been demonstrated (Suppl. Fig. 4B) and also in T98G cells [20,21] and glioma explants [22]. Here, we investigated the *in vivo* effect of two different doses (0.5 mg/kg and 1.5 mg/kg) of Bortezomib (BOR) on tumor growth when administered at the beginning of the day in the light phase or at night in the dark (Fig. 5). Controls were treated using DMSO as vehicle at the same time during the day or night. At the highest dose used (1.5 mg/kg), the drug completely inhibited tumor growth at both treatment times (day and night) (Fig. 5, dashed lines) with 97 % and 102 % of tumor growth inhibition (TGI) after day and night treatment, respectively, as determined by Rios-Doria et al [23] with modifications (Table 1). A side effect of the high dose (1.5 mg/kg) of BOR on the animals was a 12-20% weight loss along the days of treatment (Suppl. Fig. 8, dash lines); the low dose (0.5 mg/kg) did not cause any significant weight loss along the days of treatments for either condition (Suppl. Fig. 8 continuous red and blue lines). At the low dose of BOR, the nocturnal treatment (continuous blue line) showed a marked effect on tumor volume compared with daytime treatment (continuous red line) (Fig. 5B) with a TGI of 70% for the night treatment and only 18% for the day treatment (Table 1). Post hoc comparison showed that tumor volume values in animals treated with the high concentration (1.5 mg/kg) of BOR at the beginning of the day were significantly lower than those in BOR-(0.5 mg/Kg) and DMSO-treated animals after 8 days of treatment (*p < 0.05 as determined by two-way ANOVA with Bonferroni post hoc test). Although there was no difference in tumor volume between the two concentrations for the night treatment, the volumes were significantly lower than those in DMSO-treated animals after 8 days of treatment (*p < 0.05 as determined by two-way ANOVA with Bonferroni post hoc test). Results clearly show that both BOR doses used at night were effective and that furthermore, the lower dose, a third of the high dose, caused no significant side effects in terms of weight loss.

**Table 1:** Tumor Growth Inhibition (TGI). Tumor growth inhibition (%) was calculated according to Rios et al (2020) with modifications, as follows: 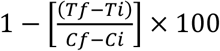, where *Tf* is the tumor volume of BOR-treated mice at day 8 of treatment, *Ti* is the tumor volume of BOR-treated mice at day 1 of treatment, *Cf* is the tumor volume of control group at day 8 of treatment and *Ci* is the tumor volume of control group at day 1 of treatment. See text for further detail. BOR = Bortezomib.

|  | Tumor growth inhibition (TGI, %) |  |
| --- | --- | --- |
|  | Day treatment | Night treatment |
| Low BOR dose (0.5 mg/kg) | 18 | 70 |
| High BOR dose (1.5 mg/kg) | 97 | 102 |

**Fig. 5:**
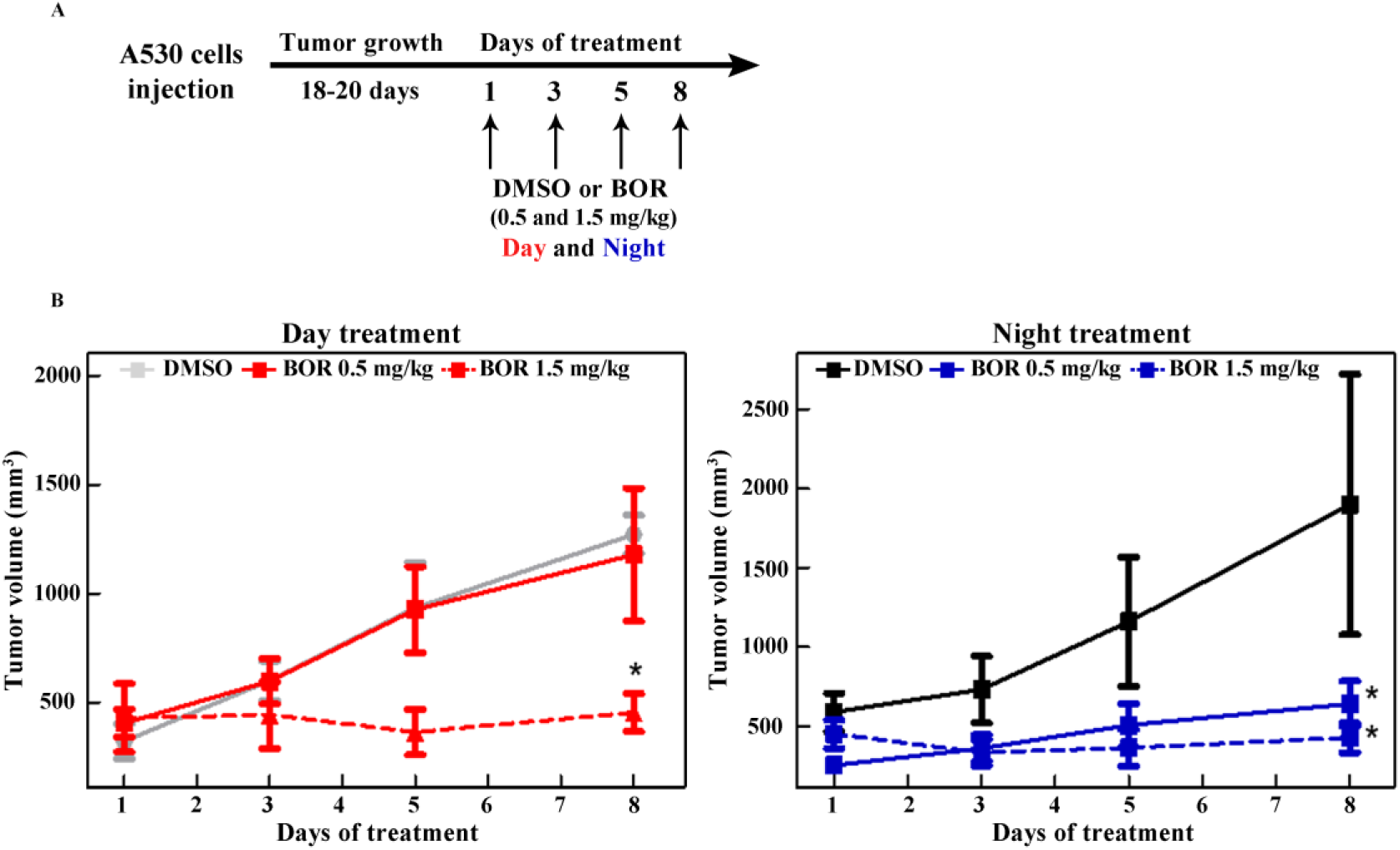
Day/night differences in the susceptibility to Bortezomib (BOR) treatment. **A**. Chemotherapeutic treatment scheme with BOR or DMSO (vehicle) for tumor-bearing C57BL/6 mice after injection of A530 glioma cells. Synchronized A530 cells (1 × 10^5^ cells in 10 µl of PBS) were injected into the sciatic nerve zone of C57BL/6 mice (A530 cells injection) and 18-20 days post-injection, mice were randomly separated into 6 groups for treatment with BOR (0.5 or 1.5 mg/kg) or DMSO (vehicle) at early day (red) or night (blue) for 8 days as indicated by the arrows; tumor volume was periodically estimated as indicated in Methods. **B**. Tumor volume growth as a function of days of treatment with BOR (0.5 or 1.5 mg/kg) or DMSO after day (left panel) or night (right panel) administration for 8 days. Day treatment (red) is shown in the left panel. Results show that the tumor volume of mice treated with a high dose of BOR (1.5 mg/kg, dash red line) significantly differs from that in animals treated with the low dose of BOR (0.5 mg/kg, red line) or DMSO (vehicle, gray line) at day 8 post-treatment (*p < 0.05). Night treatment (blue) is shown in the right panel. Results demonstrate that tumor volume of mice treated with low (0.5 mg/kg, blue line) or high (1.5 mg/kg, dash blue line) doses of BOR significantly differs from that in animals treated with DMSO (vehicle, black line) at day 8 post-treatment (*p < 0.05). Results are mean ± SEM of four independent experiments; n = 3-5 mice per group. *p < 0.05 by two-way ANOVA and Bonferroni multiple comparison test. See text for further detail.

## Discussion

Results presented herein clearly show the existence of a precise time-related control of tumor formation and growth. Notably, the state and susceptibility of the host to a cancer challenge appeared to be critical at times of higher levels of activity and metabolism. Being nocturnal mammals, mouse activity and metabolism are high at night. In the present work, we evaluated whether there was a temporary regulation of tumor growth in an *in vivo* murine model, using A530 glioma cells derived from the peripheral nervous system and B16 melanoma cells; both cells types are capable of generating tumors once injected into C57BL/6 mice during the day or night.

An important body of evidences strongly supports the existence of precise cross-talk between cancer processes and the circadian clock [3,4]. Moreover, under certain conditions, the escape from circadian physiology is crucial to tumorigenesis; in fact, the homeostasis of cells and tissues can be bypassed under the neoplastic condition to achieve rapid and exacerbated proliferation, a significant increase in metabolic requirements, immune evasion and resistance to apoptosis, and the generation of a convenient inflammatory environment highly favorable for tumor growth, all identified as cancer hallmarks [1].

We first investigated the oscillatory capability of A530 tumor cells in culture after synchronization. Cells displayed a significant temporal variation in the expression of clock genes (*Bmal*1, *Per*1), clock-controlled genes (*Chokα*) and lipid synthesizing enzymes genes (*Lipin1* and *Pcyt2*) and in the redox state, as well as in the susceptibility to chemotherapeutic treatment with BOR over time (Suppl. Figs. 2-4). These observations agree with previous reports obtained with other tumor cells in culture such as T98G and HepG2 cells derived from human glioblastoma and hepatocarcinoma, respectively [20,21]. Moreover, when comparing these results with those obtained for the T98G cell line, we may infer that A530 cells in culture contain a weak or damped oscillator with a ∼12-18 h period. One possible explanation for this damping behavior could be that A530 cells may require a stronger synchronizing cue such as a highly concentrated serum shock or a pulse of dexamethasone rather than a simple culture medium exchange to strengthen their circadian clock function. On the other hand, it should be noted that T98G cells deriving from human glioblastoma are not capable of generating tumors in C57BL/6 mice; thus, for the *in vivo* experiments, the tumor cells of choice were murine A530 and B16 cells.

### Regulation of the circadian system in tumor growth

The balance between the organism and ambient LD cycles maintains organism homeostasis. Studies in animal models show that clock disruption due to changes in the LD or feeding/fasting cycles, or to genetic mutations of clock genes, induce pathological changes that significantly affect human and animal health [3,4,12,24–26]. In fact, disruption of circadian rhythms may play a fundamental role in tumorigenesis and enable the establishment of cancer hallmarks. In this respect, taking into consideration the high cancer risk reported among shift workers, the World Health Organization (WHO) through the International Agency for Research on Cancer (IARC) has considered circadian disruption to be a potential human carcinogen of type 2A [27].

Because of several processes that drive tumorigenesis, known as cancer hallmarks, it can be inferred that escape from circadian regulation may help facilitate the transformation of normal to malignant cells through sustained proliferative signaling, genome instability and mutation, deregulation of cellular metabolism, tumor promotion of inflammation, and immune destruction avoidance. With respect to sustaining proliferative signaling, DNA replication in many organisms takes place mostly at night [28–31], encouraging cells to divide within a precise time-window and thus restricting uncontrolled division.

While the host’s differential time-of-day responses to lethal infections and endotoxins were first demonstrated almost 60 years ago [32], recent studies clarified that many aspects of immune functions are also under the control of the circadian clock [33,34]. These include immune cell trafficking, host-pathogen interactions and activation of innate and adaptive immunity [35–37]. Thus, it is suggested that circadian signals are involved in controlling the magnitude of the immune response in a time-dependent manner. In this context, our findings are novel and strengthen the link between the moment of the challenge and the triggered response. In the present paper we have shown that the injection of tumor cells with the consequent formation of tumors is subject to precise temporal control. Thus, tumors in animals injected at the beginning of the night have a faster growth rate than those injected at the beginning of the day, with significantly less duplication time as indexed by DT (Fig. 1). These results were observed with two cell models, A530 tumor cells of the peripheral nervous system and B16 melanoma cells (Fig. 1, 2), suggesting that it could be a universal phenomenon in time-regulated tumor formation and development. Moreover, the experiments carried out in constant darkness demonstrate the existence of a true endogenous rhythm in the regulation of tumor growth: it was observed that the day/night differences were also found between the subjective day and night in the absence of light signals after 72 h in DD. Our findings suggest temporary regulation essentially by the host, since no significant differences were found in the tumor growth rate when the A530 cells were synchronized at different times in culture and then injected at the same time in the mice.

In terms of day/night changes, the immune system is strongly influenced by time-of-day cues both under physiological conditions or in response to a diversity of external and pathogenic challenges [33] and involves different populations of immune cells and signaling molecules (cytokine and chemokines) along day and night. The disruption of circadian rhythms promotes the attenuation of activity by natural killer (NK) and natural killer T (NKT) cells, facilitating lung tumor progression [38]. In addition, it was shown that the initial hours in the generation of the adaptive response and the number of cells present in lymph nodes could be critical in the regulation of the triggered response (Moon et al., 2007). Moreover, susceptibility to endotoxic shock induced by lipopolysaccharide (LPS) in mice is time-dependent in a regular LD cycle and linked to time-dependent patterns in cytokine secretion [39]. A lethal administration of LPS reduces survival when administered at night compared to animals challenged in the morning [40]. In this context, rodents subjected to experimental jet-lag exhibited a marked alteration in rhythms of cytolytic factors and cytokines. These observations in combination with our findings, strengthen the importance of the state of the host and the timing of the challenge in triggering potential responses.

Some studies indicate that clock genes could act as tumor suppressors [41] while other findings suggest a more complex relationship between circadian rhythms and cancer [42]. Puram and collaborators reported that a disruption of the circadian circuit alters the growth of murine leukemia cells *in vivo* and human cell lines in culture, inducing myeloid differentiation that reduces the expression of the c-kit surface marker, a typical cell marker of leukemia [43]. They also showed that CLOCK and BMAL1 regulate cell proliferation and cell cycle progression in acute myeloid leukemia cells, consistent with phenotypes reported in other models [28,44,45]. On the contrary, Jiang and colleagues have proposed BMAL1 as a possible tumor suppressor gene in pancreatic cancer by positively regulating the p53 pathway of tumor suppression which inhibits the formation and invasion of pancreatic cancer [46]. In the tongue squamous cell carcinoma, it was shown that ectopic expression of BMAL1 inhibits cell proliferation, migration and invasion *in vitro*, and tumor growth in mouse models. In turn, over-expressing BMAL1 cells showed a relative increase in apoptosis after exposure to paclitaxel and increased susceptibility to the drug in the *in vivo* models [47]. Our results add to this evidence since we observed a higher growth rate in tumors injected with A530 A5 cells that expressed lower levels of *Bmal1* having an altered molecular clock (Fig. 3). In turn, A530 A5 cells in culture showed a higher proliferative state evidenced by a higher proportion of cells in the S-G_2_ / M phase of the cell cycle. These observations suggest the participation of the clock in acquiring the cancer hallmark related to sustained proliferation. Considering the key role that BMAL1 plays in the transduction of intracellular signals and regulation of different cellular processes, such as cell cycle progression, lipid and glucose metabolism, redox reactions and stress response [48–50], we suggest that a decrease in the expression of BMAL1 favors tumor growth as shown in Fig. 3.

In terms of gene expression, a significant altered expression of clock and clock-controlled genes was observed in tumor samples compared with surrounding tissues as found in *Per*1 and *Chokα* (Fig. 4). In this respect, there is growing evidence that circadian disruption in gene expression may alter risk and aggressiveness of tumors such as glioblastoma and other classes: *Per*1/2 expression was shown to be decreased or altered in cancer cells [4,51,52] while *Chokα* was shown to be regulated in a circadian manner in normal hepatic tissue [44,53,54] with exacerbated expression in some tumor cells [55].

In relation to the chemotherapeutic treatment with BOR, our results show that when a low dose was utilized, at which no major side effects in body weight were observed, the nocturnal administration was remarkably more effective than when the same drug concentration was administered during the day (Fig. 5). The nocturnal administration caused tumor growth inhibition close to 70% as compared with only 18% of inhibition during the day. This demonstrates that there is a tight daily regulation of BOR chemotherapy outcome with higher efficacy at night. Observations may reflect circadian variations in drug availability, activity and metabolism as well as in xenobiotic detoxification enzymes, and these variations may also cooperate to maximize therapeutic effects and minimize side effects, especially toxicity in non-cancerous cells [4]. Furthermore, diverse chemotherapeutic agents acting at different cellular levels and mechanisms can be firmly modulated by the circadian clock at specific drug concentrations in the procedure so-called “chronotherapy” [56]; some of them act on the proteasome (BOR) as shown herein, others induce DNA damage (temozolamide and cisplastin) [22,57] while REV-ERB agonists such as the pyrrole derivative SR9009 and others act on the clock-related cellular metabolism [58]. Nevertheless, most of them display circadian variations in cell susceptibility to cytotoxicity with precise temporal windows exhibiting the highest antitumor drug effect. In some cases, when different drugs were used in combination, low drug doses of each seemed to have potent synergic effects, as was recently reported [20]. Tumor cell-intrinsic circadian rhythms are present in GBM tumors and can regulate DNA alkylator temozolamide cytotoxicity [22]. Moreover, the action of cisplatin that kills cancer cells by damaging their DNA is controlled by two different circadian programs in mouse tissues [57]. Our results with BOR show that night administration offers a temporal window exhibiting higher drug efficacy when mice are active in their metabolism, locomotor activity and feeding habits. Although animals were more susceptible to the tumor cell challenge and exhibited a higher tumor growth rate at night, they also responded more efficiently to chemotherapy at this time, and especially at lower doses with no undesired side effects. The integration of circadian biology into cancer research offers new possibilities to improve cancer treatments and make them more effective, potentially translating into a better health outcome for the patients.

### Concluding remarks

The *in vivo* experiments strongly support the idea that the circadian clock is closely linked to the growth of tumors generated by the injection of cancer cells and their chemotherapeutic treatment. Indeed, our findings clearly show a clock-controlled process specific to the host, exhibiting a precise time-control of cancer progression with higher rates of tumor growth and chemotherapy susceptibility at night. Moreover, after knocking down the expression of the clock gene *Bmal1*, higher tumor growth rates were observed than in tumors generated from control cells, strengthening the relationship between the transcriptional machinery of the circadian clock and tumor development. Understanding and delving into tumor regulation from a chronobiological viewpoint will help in the design of new treatments that maximize therapeutic benefits at certain times of the day while minimizing the adverse effects of therapy.

## Materials and Methods

### Cell Culture and synchronization

**A530 cells** were isolated from malignant peripheral nerve sheath tumor (MPNST) of NPcis (*Trp53*^*+/-*^; *Nf1*^*+/-*^) mice in a background C57BL/6, an animal model for the human neurofibromatosis type I (Suppl. Fig. 1). NPcis mice bare a disrupted allele for both the *Trp*53 and the *Nf*1 tumor suppressor genes; the loss of heterozygosis determines the spontaneous development of central and peripheral nervous system tumors by the age of 6-7 month with a 100% penetrance [17]. Peripheral nervous tumor was dissociated and p75^+^PI^-^ cells separated using flow cytometry cell sorting (Suppl. Fig. 1A-B). p75^+^ cells have been found to be the most abundant proliferating cells in neurofibromas [59] with cancer stem cell properties [60]. Tumor tissue was dissociated for 30 minutes at 37°C in 1 mg/mL type IV collagenase followed by 20 minutes at 37°C in 0.5% trypsin/EDTA. The reaction was quenched with DMEM/F12 plus 20% fetal bovine serum (FBS), 1% penicillin/streptomycin and 25mg/mL deoxiribonuclease type I (DNase I). Cells were centrifuged, resuspended in DMEM/F12 plus 20% FBS and 1% penicillin/streptomycin, triturate and filtered through nylon screen (45 mm) to remove aggregates. P75^+^ PI^-^ cells were isolated by cell sorting after incubation with antibody against p75 and propidium iodide (PI) staining. Cells were tested positive for glial cell markers and negative for mycoplasma contamination (Suppl. Fig. 1D). Cells were grown in Dulbecco modified Eagles low glucose medium (DMEM, Gibco) and Neurobasal medium (Gibco) supplemented with 20% FBS (Gibco), FGF (20 ng/ml), EGF (20 ng/ml), β-mercaptoethanol (55 µM) and retinoic acid (0.12 µM) an incubated in a CO_2_ incubator at 37°C for 24-48 h. Cells were synchronized by culture medium exchange and allowed to grow in complete medium. For studies in culture, after synchronization (time 0), cells were collected at 6 h intervals during 36 h.

**B16 cells** are derived from the mouse melanoma of skin (ATCC^®^ CRL-6322^(tm)^, Manassas, VA, USA) and tested negative for mycoplasma contamination. Cells were grown in DMEM supplemented with 10% FBS. Previous to the injection in mice, B16 cells were synchronized by culture medium exchange and allow growing in complete medium for 24 h.

### Characterization of A530 cells in culture

Cells isolated from MPNSTs were named A530 cells (Suppl. Fig. 1A). These cells can grow in suspension without contact inhibition and are able to form spheres and lumps as described for typical malignant cells (Suppl. Fig 1B). Genomic analysis of *Nf*1 and *Trp*53 genes from A530 cells and C57BL/6 mice DNA evidenced the loss of both wild type (WT) copies of these genes in A530 cells (Suppl. Fig. 1C). Moreover, A530 cells were characterized as glioma cells that expressed typical glial markers such as glutamine synthase (GS), glial fibrillary acidic protein (GFAP) and vimentin, as well as stem cell markers such as SOX2, a persistent marker for multipotent neural stem cells derived from embryonic stem cells, and S100, a family of proteins normally present in cells derived from the neural crest (Schwann cells, and Schwann cells) and other cells (Suppl. Fig. 1D).

### Redox State in A530 Cells-Temporal Regulation

To investigate the metabolic/redox clock in A530 cells, the redox state was assessed in cultured cells after culture medium exchange synchronization. For this, A530 cells grown as indicated above were harvested at 6 h intervals after synchronization during 36 h and the redox state of the cells was analyzed by flow cytometry as described [20, 21, 61]. Briefly, the growth medium was removed, cells were washed with cold PBS 1X and harvested by trypsinization. 4 × 10^5^ cells were resuspended in phosphate buffer saline (PBS) 1X and incubated with 2′, 7′-Dichlorodihydrofluorescein diacetate at 2 μM final concentration for 40 min at 37°C. The cells were washed twice with PBS 1X and the fluorescence intensity was measured by flow cytometry at 530 nm when the sample was excited at 485 nm. A negative control including cells without the fluorescent probe was used. When the fluorescence intensity was analyzed, a significant temporal variation in the cellular redox state was observed (Suppl. Fig. 2A) (p < 0.04 by Kruskal-Wallis) with higher ROS levels at 0-6 h, a decrease by time 12, and then a damped oscillation was observed.

### Bortezomib Treatment of Synchronized A530 Cell Cultures and Determination of Viability

Cells were plated in 96-well plates at a density of 1×10^4^ and were allowed to attach overnight at 37°C. Cultured cells were synchronized by culture medium exchange and then either left untreated or treated with Bortezomib (BOR) at different times (h) at 500 nM final concentration for 24 h. After incubation, 10 μl of MTT reagent (5 mg/ml; Sigma) were added to each well, and plates were further incubated for 2 h at 37°C as described [21, 62, 63]. Then, 100μl of DMSO: isopropanol (1:1, v/v) was added to each well followed by incubation for a few min at room temperature protected from light. Samples were analyzed at a wavelength of 570 nm with a reference at 650 nm using an Epoch Microplate Spectrophotometer. Results revealed a significant temporal effect of the drug treatment (p < 0.006 by Kruskal-Wallis), with the lowest levels of viability at 0, 18 and 36 h (Suppl. Fig. 2B). It is noteworthy that fluctuations observed seemed to display a 12-18 h short period during a first oscillation which damped at longer times likely reflecting either a weak *in vitro* oscillator or a not strong enough synchronizing signal.

### RNA isolation and Reverse Transcription

Total RNA was extracted from cell homogenates using TRIzol® reagent following the manufacturer’s specifications (Invitrogen). The yield and purity of RNA were estimated by optical density at 260/280 nm. 2 µg of total RNA was treated with DNAse (Promega) and utilized as a template for the cDNA synthesis reaction using MMLV reverse transcriptase (Promega) and an equimolar mix of random hexamers (Biodynamics) in a final volume of 25 µL according to the manufacturer’s indications.

### Real-time PCR (qPCR)

Quantitative PCR was performed using SYBR Green or TaqMan Gene Expression Assays in a Rotor Q Gene (QIAGEN). The primer/probe sequences are summarized in Suppl. Table 1. The amplification mix contained 2 µL of the cDNA, 2 µL 20X mix primer/probe or 300 nM Forward-Reverse TBP primers, and 7.5 µL of Master Mix 2X (Applied Biosystems) in a final volume of 15 µL. The cycling conditions were 10 min at 95.0°C, and 45 cycles of 95.0°C for 15 sec, 60.0°C for 30 sec and 72°C for 30 sec. The standard curve linearity and PCR efficiency (E) were optimized. We used the 2 ^-ΔΔCT^ according to Livak and Schmittgen [64] and Larionov et al. [65] and *tbp* as the reference gene [54]. Each RT-PCR quantification experiment was performed at least in duplicate (TaqMan or SYBR) for each sample (n=2-5/sample).

### Determination of Gene Expression in synchronized A530 cell cultures

A530 cells were grown in culture and synchronized by a culture medium exchange (time 0). After synchronization, samples were collected at different times and RNA extracted and processed as described above. Cells displayed temporal variations in the expression of clock genes (*Bmal*1, *Per*1) and genes encoding for phospholipid synthesizing enzymes such as *Lipin1, ChoKα and Pcyt*2 (Suppl. Fig. 3). *Lipin1* encodes for the phosphatidate phosphohydrolase 1 (PAP1), *ChoKα* for a choline kinase isoform of the phosphatidylcholine synthesis (Kennedy pathway) and *Pcyt*2 gene encodes for the ethanolamine phosphate citidyltransferase enzyme (Kennedy pathway). Expression of mRNAs exhibited a significant effect of time (p < 0.04 by Kruskal-Wallis) with different phases and peaking around 20 and 40 h post-synchronization.

### Genomic analysis of *Nf*1 and *Trp*53 genes from A530 cells

Genomic DNA was extracted from homogenates using TRIzol® reagent following the manufacturer’s specifications (Invitrogen). The yield and purity of DNA were estimated by optical density at 260/280 nm. Briefly, DNA was precipitated with 100% ethanol after RNA isolation from homogenates. The DNA pellet was washed twice with sodium citrate in 10% ethanol and resuspended in nuclease-free water. 0.25 µg of DNA was used to amplify specific regions of *Trp*53 and *Nf*1 gene using Go-Taq DNA Polymerase (Promega). The primers sequences are summarized in Suppl. Table 2. The cycling conditions were 3 min at 94°C, and 35 cycles of 94 °C for 30 sec, 54 °C for 30 sec and 72°C for 60 sec. Amplicons were separated on an agarose gel by electrophoresis and visualized by ethidium bromide staining on a transilluminator. Results are shown in Suppl. Fig. 1C.

### Propidium Iodide Staining and Flow cytometry

A530 cells were grown in complete culture medium as described above for 24-48 h and then synchronized by culture medium exchange. Cells were harvested 24 h post-synchronization, washed with cold PBS 1X, and fixed with cold 70% (v/v) ethanol for at least 24 h at -20°C. Cell pellets were resuspended in 150 µL of staining solution (PBS containing 50 μg/mL propidium iodide and 10 μg RNAse A) as reported [54]. Cell cycle analysis was performed using 50,000 cells on a flow cytometer (DB Bioscience). FlowJo software was used for the analysis.

### Knock-down of *Bmal1* expression in A530 cells by CRISPR/cas9 technology

*Bmal1* expression was disrupted in transfected A530 cells using the CRISPR/Cas9 genomic editing tool as previously described [21, 66]. Briefly, we designed single guide RNAs specifically targeting exon 1 of the mouse *Bmal1* gene and subcloned it into the PX459 vector (Addgene) to obtain the PX459-*Bmal1* plasmid. The primer sequence corresponding to the single guide RNA was 5’ AGGTGCCTGTTTACCCGCGC 3’ and the complementary sequence 5’TCCACGGACAAATGGGCGCG 3’. A530 cells were transfected with Lipofectamine 2000 (Invitrogen) and selected with puromycin (2 μg/ml) for 5 days. The pool of selected A530 cells was named as A530 A5. The disruption of *Bmal1* gene expression was checked by RT-PCR and shown to be at least 50% lower than in A530 cells. Additional effects were found on the cell cycle phases by flow cytometry and on the expression of the downstream target gene *Per1* by RT-PCR. Results are shown in Suppl. Fig. 7.

### Immunocytochemistry (ICC)

ICC was performed as described [67]. Briefly, cultured cells were fixed for 15 min in 4% paraformaldehyde in PBS and 10 min in methanol. Coverslips were washed in PBS, treated with blocking buffer (PBS supplemented with 0.1% BSA, 0.1% Tween 20 and 0.1% glycine) and incubated overnight with primary antibodies. The cells were then rinsed in PBS and incubated with secondary antibodies for 1 h at room temperature (RT). Coverslips were finally washed thoroughly and visualized by confocal microscopy (FV1200; Olympus, Tokyo, Japan). Cellular nuclei were visualized by DAPI staining. To assess PER1-like protein immunoreactivity, synchronized A530 cells were treated as described above for ICC with the specific primary PER1 antibody (Suppl. Table 3). Cells displayed a temporal variation in PER1-like protein levels along 30 h with the highest values at a time window ranging from 12-24 h and lowest levels at 2, 6 and 30 h post synchronization (p < 0.0001 by Kruskal-Wallis) (Suppl. Fig. 4).

### Animal Handling

C57BL/6 mice (8-12 weeks of age) were housed under a regular 12-hour light/dark (LD) cycle (lights-on at 7:00 AM and lights-off at 7:00 PM) with food and water *ad libitum*. Time expressed as “Zeitgeber Time” (ZT; where lights were turned on at ZT 0 and off at ZT 12). To assess free-running experiments in constant darkness (DD), mice were entrained to a regular LD cycle for 7 days and then released to DD for 48-72 h. Because mice have free-running periods very close to 24 h and they would not have shifted significantly after 48-72 h of DD, times of tumor cells injection are designated with respect to the previous entraining LD cycle (or zeitgeber). Thus, ZT 0 corresponds to the phase of the previous dark-light transition (subjective dawn), while ZT 12 corresponds to the time of the light-dark transition (subjective dusk). After the injection of tumor cells, animals were maintained on DD for another 15 days. All animal procedures were performed following the protocol approved by Guide to the Care and Use of Experimental Animals, published by the Canadian Council on Animal Care and approved by the local animal care committee (School of Chemistry, National University of Cordoba; Exp.15-99-39796) and CICUAL (Institutional Committee for the Care and Use of Experimental Animals Exp. Nr. 2139/2017).

### Tumor growth experiments

For A530 cells injection, cells synchronized by culture medium exchange were harvested by trypsinization and washed with cold PBS 1X. 1 × 10^5^ cells resuspended in 10 µl of PBS were injected into the sciatic nerve zone of C57BL/6 mice at the beginning of the day (ZT 1-2) or night (ZT 12-13). Tumors became palpable around 15 days post-injection. Tumor growth was measured periodically with calipers and tumor volumes were estimated according to the formula: (long dimension) * (short dimension) * (height dimension). Mice were sacrificed around day 28-30 post-inoculation (Suppl. Figure 5). For B16 cells injection, 5 × 10^5^ cells resuspended in 50 µl of PBS were injected subcutaneously into the right flank of C57BL/6 mice at the beginning of the day or night. Tumors became palpable around 7 days post-injection and tumor growth was measured periodically using a caliper as described above. Tumor volumes were calculated according to the formula: [(width)^2^ × (length)]/2. Mice were sacrificed around 15 days post-injection.

Individual tumor growth rates were calculated by linear regression of log transformed tumor volumes of each group according to [68]. At early-stages of tumor growth, this exponential model supposes that cells divide regularly, creating two daughter cells each time [69], in which growth is proportional to the population. However, this model may fail at later stages when angiogenesis and nutrient depletion begin to play a role [70, 71].

**Doubling time (DT):** Tumor growth can be also characterized by the tumor volume doubling time (DT). This parameter was determined as follows: DT = In2/[In (Vf/Vi)/(t_f_-t_i_)], where Vf and Vi represent the tumor volume measured at the end (t_f_) and beginning (t_i_) of the experiment, respectively [72, 73].

**Relative tumor volume** was calculated according to the formula: Vx/Vi, where Vx and Vi represent the tumor volume measured at day “x” and at the beginning of the experiment, respectively.

### *In vivo* Bortezomib treatment

For Bortezomib (BOR) chronotherapy experiments, C57BL/6 mice were injected with A530 cells into the sciatic nerve zone as described above. After 18-20 days post-injection, mice were randomly separated into 6 groups. The experimental protocol shown in Fig. 5A consisted of 4 doses of BOR at the concentration of 0.5 or 1.5 mg/kg resuspended in 30 μL of DMSO, for one week administered intraperitoneal at the beginning of the day or night. Control groups were injected with DMSO (vehicle). Animal weight and tumor volume were periodically measured as described above. To evaluate BOR treatment efficacy, tumor growth inhibition (TGI %) was calculated according to Rios et al. [23] with modifications as follows: 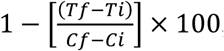, where *Tf* is the tumor volume of BOR-treated mice at day 8 of treatment, *Ti* is the tumor volume of BOR-treated mice at day 1 of treatment while *Cf* is the tumor volume of control group (DMSO) at day 8 of treatment and *Ci* is the tumor volume of control group at day 1 of treatment.

### Statistics

Statistical analyses involved a one-or two-way analysis of variance (ANOVA) to test the time or drug treatment effects and Kruskal-Wallis when the normality of residuals was infringed. Pairwise comparisons were performed by the Duncan and Bonferroni test when appropriate. The Student t-test (one-tailed) was used to compare the differences between the means of tumor growth in each experiment, and a non-parametric t-test Mann-Whitney when the model assumptions were infringed. In all cases, significance was considered at p < 0.05. All statistical analysis was performed with Prism5 GraphPad Software.

## Acknowledgments

This work has been supported by Agencia Nacional de Promoción Científica y Técnica (FONCyT, PICT 2016 N° 0187 and 2017 N° 631), Consejo Nacional de Investigaciones Científicas y Tecnológicas de la República Argentina (CONICET) (PIP 2014) and Secretaría de Ciencia y Tecnología de la Universidad Nacional de Córdoba (SeCyT-UNC, Consolidar 2018-2022).

## Notes

### Competing Interest Statement

The authors have declared no competing interest.

## References

1. Hanahan D, Weinberg RA. Hallmarks of Cancer: The Next Generation. Cell. 2011;144: 646–674. doi:10.1016/j.cell.2011.02.013

2. Schernhammer ES, Laden F, Speizer FE, Willett WC, Hunter DJ, Kawachi I, et al. Rotating Night Shifts and Risk of Breast Cancer in Women Participating in the Nurses’ Health Study. JNCI J Natl Cancer Inst. 2001;93: 1563–1568. doi:10.1093/jnci/93.20.1563

3. Lahti T, Merikanto I, Partonen T. Circadian clock disruptions and the risk of cancer. Ann Med. 2012;44: 847–853. doi:10.3109/07853890.2012.727018

4. Sulli G, Lam MTY, Panda S. Interplay between Circadian Clock and Cancer: New Frontiers for Cancer Treatment. Trends in Cancer. Cell Press; 2019. pp. 475–494. doi:10.1016/j.trecan.2019.07.002

5. Mohawk JA, Green CB, Takahashi JS. Central and peripheral circadian clocks in mammals. Annu Rev Neurosci. 2012/04/05. 2012;35: 445–462. doi:10.1146/annurev-neuro-060909-153128

6. Takahashi JS. Molecular components of the circadian clock in mammals. Diabetes Obes Metab. 2015;17 Suppl 1: 6–11. doi:10.1111/dom.12514

7. Preitner N, Damiola F, Lopez-Molina L, Zakany J, Duboule D, Albrecht U, et al. The orphan nuclear receptor REV-ERBalpha controls circadian transcription within the positive limb of the mammalian circadian oscillator. Cell. 2002;110: 251–60. Available: http://www.ncbi.nlm.nih.gov/pubmed/12150932

8. Sato TK, Panda S, Miraglia LJ, Reyes TM, Rudic RD, McNamara P, et al. A Functional Genomics Strategy Reveals Rora as a Component of the Mammalian Circadian Clock. Neuron. 2004;43: 527–537. doi:10.1016/j.neuron.2004.07.018

9. Bollinger T, Schibler U. Circadian rhythms – from genes to physiology and disease. Swiss Med Wkly. 2014;144: w13984. doi:10.4414/smw.2014.13984

10. Dibner C, Schibler U. Circadian timing of metabolism in animal models and humans. J Intern Med. 2015;277: 513–527. doi:10.1111/joim.12347

11. Gooley JJ, Chua EC-P. Diurnal Regulation of Lipid Metabolism and Applications of Circadian Lipidomics. J Genet Genomics. 2014;41: 231–250. doi:10.1016/j.jgg.2014.04.001

12. Fu L, Kettner NM. The circadian clock in cancer development and therapy. Progress in Molecular Biology and Translational Science. Elsevier B.V.; 2013. pp. 221–282. doi:10.1016/B978-0-12-396971-2.00009-9

13. Takahashi JS, Hong H-K, Ko CH, McDearmon EL. The genetics of mammalian circadian order and disorder: implications for physiology and disease. Nat Rev Genet. 2008;9: 764–75. doi:10.1038/nrg2430

14. Wesseling P, Capper D. WHO 2016 Classification of gliomas. Neuropathol Appl Neurobiol. 2018;44: 139–150. doi:10.1111/nan.12432

15. Lenting K, Verhaak R, ter Laan M, Wesseling P, Leenders W. Glioma: experimental models and reality. Acta Neuropathologica. Springer Verlag; 2017. pp. 263–282. doi:10.1007/s00401-017-1671-4

16. Stylli SS, Luwor RB, Ware TMB, Tan F, Kaye AH. Mouse models of glioma. Journal of Clinical Neuroscience. Churchill Livingstone; 2015. pp. 619–626. doi:10.1016/j.jocn.2014.10.013

17. Reilly KM, Loisel DA, Bronson RT, McLaughlin ME, Jacks T. Nf1;Trp53 mutant mice develop glioblastoma with evidence of strain-specific effects. Nat Genet. 2000;26: 109–113. doi:10.1038/79075

18. Chen D, Frezza M, Schmitt S, Kanwar J, P. Dou Q. Bortezomib as the First Proteasome Inhibitor Anticancer Drug: Current Status and Future Perspectives. Curr Cancer Drug Targets. 2011;11: 239–253. doi:10.2174/156800911794519752

19. Mehrara E, Forssell-Aronsson E, Ahlman H, Bernhardt P. Quantitative analysis of tumor growth rate and changes in tumor marker level: Specific growth rate versus doubling time. Acta Oncol (Madr). 2009;48: 591–597. doi:10.1080/02841860802616736

20. Wagner PM, Monjes NM, Guido ME. Chemotherapeutic Effect of SR9009, a REV-ERB Agonist, on the Human Glioblastoma T98G Cells. ASN Neuro. 2019;11: 175909141989271. doi:10.1177/1759091419892713

21. Wagner PM, Sosa Alderete LG, Gorné LD, Gaveglio V, Salvador G, Pasquaré S, et al. Proliferative Glioblastoma Cancer Cells Exhibit Persisting Temporal Control of Metabolism and Display Differential Temporal Drug Susceptibility in Chemotherapy. Mol Neurobiol. 2018 [cited 8 Feb 2019]. doi:10.1007/s12035-018-1152-3

22. Slat EA, Sponagel J, Marpegan L, Simon T, Kfoury N, Kim A, et al. Cell-intrinsic, Bmal1-dependent Circadian Regulation of Temozolomide Sensitivity in Glioblastoma. J Biol Rhythms. 2017;32: 121–129. doi:10.1177/0748730417696788

23. Rios-Doria J, Stevens C, Maddage C, Lasky K, Koblish HK. Characterization of human cancer xenografts in humanized mice. J Immunother Cancer. 2020;8: e000416. doi:10.1136/jitc-2019-000416

24. Lahti TA, Partonen T, Kyyrönen P, Kauppinen T, Pukkala E. Night-time work predisposes to non-Hodgkin lymphoma. Int J Cancer. 2008;123: 2148–2151. doi:10.1002/ijc.23566

25. Takahashi JS, Hong H-K, Ko CH, McDearmon EL. The genetics of mammalian circadian order and disorder: implications for physiology and disease. Nat Rev Genet. 2008;9: 764–775. doi:10.1038/nrg2430

26. Acosta-Rodríguez VA, de Groot MHM, Rijo-Ferreira F, Green CB, Takahashi JS. Mice under Caloric Restriction Self-Impose a Temporal Restriction of Food Intake as Revealed by an Automated Feeder System. Cell Metab. 2017;26: 267-277.e2. doi:10.1016/j.cmet.2017.06.007

27. Straif K, Baan R, Grosse Y, Secretan B, Ghissassi F El, Bouvard V, et al. Carcinogenicity of shift-work, painting, and fire-fighting. Lancet Oncol. 2007;8: 1065–1066. doi:10.1016/S1470-2045(07)70373-X

28. Matsuo T. Control Mechanism of the Circadian Clock for Timing of Cell Division in Vivo. Science (80-). 2003;302: 255–259. doi:10.1126/science.1086271

29. Plikus M V., Vollmers C, De La Cruz D, Chaix A, Ramos R, Panda S, et al. Local circadian clock gates cell cycle progression of transient amplifying cells during regenerative hair cycling. Proc Natl Acad Sci U S A. 2013;110: E2106–E2115. doi:10.1073/pnas.1215935110

30. Bjarnason GA, Jordan R. Circadian variation of cell proliferation and cell cycle protein expression in man: clinical implications. Progress in cell cycle research. Springer, Boston, MA; 2000. pp. 193–206. doi:10.1007/978-1-4615-4253-7_17

31. Smaaland R. Circadian rhythm of cell division. Progress in cell cycle research. Springer, Boston, MA; 1996. pp. 241–266. doi:10.1007/978-1-4615-5873-6_23

32. Halberg F, Johnson EA, Brown BW, Bittner JJ. Susceptibility Rhythm to E. coli Endotoxin and Bioassay. Exp Biol Med. 1960;103: 142–144. doi:10.3181/00379727-103-25439

33. Scheiermann C, Gibbs J, Ince L, Loudon A. Clocking in to immunity. Nature Reviews Immunology. Nature Publishing Group; 2018. pp. 423–437. doi:10.1038/s41577-018-0008-4

34. Rijo-Ferreira F, Takahashi JS. Genomics of circadian rhythms in health and disease. Genome Medicine. BioMed Central Ltd.; 2019. pp. 1–16. doi:10.1186/s13073-019-0704-0

35. Scheiermann C, Kunisaki Y, Frenette PS. Circadian control of the immune system. Nat Rev Immunol. 2013;13: 190–8. doi:10.1038/nri3386

36. Curtis AM, Bellet MM, Sassone-Corsi P, O’Neill LAJ. Circadian clock proteins and immunity. Immunity. 2014;40: 178–86. doi:10.1016/j.immuni.2014.02.002

37. Labrecque N, Cermakian N. Circadian Clocks in the Immune System. J Biol Rhythms. 2015;30: 277–290. doi:10.1177/0748730415577723

38. Logan RW, Zhang C, Murugan S, O’Connell S, Levitt D, Rosenwasser AM, et al. Chronic Shift-Lag Alters the Circadian Clock of NK Cells and Promotes Lung Cancer Growth in Rats. J Immunol. 2012;188: 2583–2591. doi:10.4049/jimmunol.1102715

39. Marpegan L, Leone MJ, Katz ME, Sobrero PM, Bekinstein TA, Golombek DA. DIURNAL VARIATION IN ENDOTOXIN-INDUCED MORTALITY IN MICE: CORRELATION WITH PROINFLAMMATORY FACTORS. Chronobiol Int. 2009;26: 1430–1442. doi:10.3109/07420520903408358

40. Scheiermann C, Kunisaki Y, Lucas D, Chow A, Jang J-E, Zhang D, et al. Adrenergic nerves govern circadian leukocyte recruitment to tissues. Immunity. 2012;37: 290–301. doi:10.1016/j.immuni.2012.05.021

41. Fu L, Pelicano H, Liu J, Huang P, Lee C. The circadian gene Period2 plays an important role in tumor suppression and DNA damage response in vivo. Cell. 2002;111: 41–50. Available: http://www.ncbi.nlm.nih.gov/pubmed/12372299

42. Antoch MP, Toshkov I, Kuropatwinski KK, Jackson M. Deficiency in PER proteins has no effect on the rate of spontaneous and radiation-induced carcinogenesis. Cell Cycle. 2013;12: 3673–80. doi:10.4161/cc.26614

43. Puram R V, Kowalczyk MS, De Boer CG, Al-Shahrour F, Regev A, Ebert BL. Core Circadian Clock Genes Regulate Leukemia Stem Cells in AML In Brief Disruption of the circadian rhythm machinery in AML produces anti-leukemic effects, including differentiation and depletion of disease-propagating leukemia stem cells. Accession Numbers GSE70686. 2016 [cited 27 Jun 2019]. doi:10.1016/j.cell.2016.03.015

44. Gréchez-Cassiau A, Feillet C, Guérin S, Delaunay F. The hepatic circadian clock regulates the choline kinase α gene through the BMAL1-REV-ERBα axis. Chronobiol Int. 2015;32: 774–784. doi:10.3109/07420528.2015.1046601

45. Miller BH, McDearmon EL, Panda S, Hayes KR, Zhang J, Andrews JL, et al. Circadian and CLOCK-controlled regulation of the mouse transcriptome and cell proliferation. Proc Natl Acad Sci. 2007;104: 3342–3347. doi:10.1073/pnas.0611724104

46. Jiang W, Zhao S, Jiang X, Zhang E, Hu G, Hu B, et al. The circadian clock gene Bmal1 acts as a potential anti-oncogene in pancreatic cancer by activating the p53 tumor suppressor pathway. Cancer Lett. 2016;371: 314–325. doi:10.1016/J.CANLET.2015.12.002

47. Tang Q, Cheng B, Xie M, Chen Y, Zhao J, Zhou X, et al. Circadian Clock Gene *Bmal1* Inhibits Tumorigenesis and Increases Paclitaxel Sensitivity in Tongue Squamous Cell Carcinoma. Cancer Res. 2017;77: 532–544. doi:10.1158/0008-5472.CAN-16-1322

48. Razorenova O V. Brain and muscle ARNT-like protein BMAL1 regulates ROS homeostasis and senescence: A possible link to hypoxia-inducible factor-mediated pathway. Cell Cycle. 2012;11: 213–213. doi:10.4161/cc.11.2.18786

49. Hatanaka F, Matsubara C, Myung J, Yoritaka T, Kamimura N, Tsutsumi S, et al. Genome-wide profiling of the core clock protein BMAL1 targets reveals a strict relationship with metabolism. Mol Cell Biol. 2010/10/11. 2010;30: 5636–5648. doi:10.1128/MCB.00781-10

50. Gallego M, Virshup DM. Post-translational modifications regulate the ticking of the circadian clock. Nat Rev Mol Cell Biol. 2007;8: 139. doi:10.1038/nrm2106

51. Zhanfeng N, Yanhui L, Zhou F, Shaocai H, Guangxing L, Hechun X. Circadian genes Per1 and Per2 increase radiosensitivity of glioma in vivo. Oncotarget. 2015;6: 9951–8. doi:10.18632/oncotarget.3179

52. Madden MH, Anic GM, Thompson RC, Nabors LB, Olson JJ, Browning JE, et al. Circadian pathway genes in relation to glioma risk and outcome. Cancer Causes Control. 2014;25: 25–32. doi:10.1007/s10552-013-0305-y

53. Gorné LD, Acosta-Rodríguez VA, Pasquaré SJ, Salvador GA, Giusto NM, Guido ME. The mouse liver displays daily rhythms in the metabolism of phospholipids and in the activity of lipid synthesizing enzymes. Chronobiol Int. 2015;32: 11–26. doi:10.3109/07420528.2014.949734

54. Acosta-Rodríguez VA, Márquez S, Salvador GA, Pasquaré SJ, Gorné LD, Garbarino-Pico E, et al. Daily rhythms of glycerophospholipid synthesis in fibroblast cultures involve differential enzyme contributions. J Lipid Res. 2013;54: 1798–1811. doi:10.1194/jlr.M034264

55. Gallego-Ortega D, Gómez del Pulgar T, Valdés-Mora F, Cebrián A, Lacal JC. Involvement of human choline kinase alpha and beta in carcinogenesis: A different role in lipid metabolism and biological functions. Advances in Enzyme Regulation. 2011. pp. 183–194. doi:10.1016/j.advenzreg.2010.09.010

56. Masri S, Sassone-Corsi P. The emerging link between cancer, metabolism, and circadian rhythms. Nature Medicine. Nature Publishing Group; 2018. pp. 1795–1803. doi:10.1038/s41591-018-0271-8

57. Yang Y, Adebali O, Wu G, Selby CP, Chiou YY, Rashid N, et al. Cisplatin-DNA adduct repair of transcribed genes is controlled by two circadian programs in mouse tissues. Proc Natl Acad Sci U S A. 2018;115: E4777–E4785. doi:10.1073/pnas.1804493115

58. Solt LA, Wang Y, Banerjee S, Hughes T, Kojetin DJ, Lundasen T, et al. Regulation of circadian behaviour and metabolism by synthetic REV-ERB agonists. Nature. 2012;485: 62–8. doi:10.1038/nature11030

59. Eruslanov E, Kusmartsev S. Identification of ROS Using Oxidized DCFDA and Flow-Cytometry. Methods in molecular biology (Clifton, NJ). 2010. pp. 57–72. doi:10.1007/978-1-60761-411-1_4

60. Vlachostergios PJ, Hatzidaki E, Stathakis NE, Koukoulis GK, Papandreou CN. Bortezomib Downregulates MGMT Expression in T98G Glioblastoma Cells. Cell Mol Neurobiol. 2013;33: 313–318. doi:10.1007/s10571-013-9910-2

61. Comba A, Bonnet L V., Goitea VE, Hallak ME, Galiano MR. Arginylated Calreticulin Increases Apoptotic Response Induced by Bortezomib in Glioma Cells. Mol Neurobiol. 2019;56: 1653–1664. doi:10.1007/s12035-018-1182-x

62. Ran FA, Hsu PD, Wright J, Agarwala V, Scott DA, Zhang F. Genome engineering using the CRISPR-Cas9 system. Nat Protoc. 2013;8: 2281–2308. doi:10.1038/nprot.2013.143

63. Morera LP, Díaz NM, Guido ME. Horizontal cells expressing melanopsin x are novel photoreceptors in the avian inner retina. Proc Natl Acad Sci. 2016;113: 13215–13220. doi:10.1073/pnas.1608901113

64. Liu Y, Ludes-Meyers J, Zhang Y, Munoz-Medellin D, Kim H-T, Lu C, et al. Inhibition of AP-1 transcription factor causes blockade of multiple signal transduction pathways and inhibits breast cancer growth. Oncogene. 2002;21: 7680–7689. doi:10.1038/sj.onc.1205883

65. Mehrara E, Forssell-Aronsson E, Ahlman H, Bernhardt P. Quantitative analysis of tumor growth rate and changes in tumor marker level: Specific growth rate versus doubling time. Acta Oncol (Madr). 2009;48: 591–597. doi:10.1080/02841860802616736

66. D. Sarma H, Das T, Banerjee S, Venkatesh M, B. Vidyasagar P, P. Mishra K. Studies on Efficacy of a Novel 177Lu-Labeled Porphyrin Derivative in Regression of Tumors in Mouse Model. Curr Radiopharm. 2011;4: 150–160. doi:10.2174/1874471011104020150

67. Garbarino-Pico E, Niu S, Rollag MD, Strayer CA, Besharse JC, Green CB. Immediate early response of the circadian polyA ribonuclease nocturnin to two extracellular stimuli. [cited 27 Apr 2020]. doi:10.1261/rna.286507

68. Nakahata Y, Akashi M, Trcka D, Yasuda A, Takumi T. The in vitro real-time oscillation monitoring system identifies potential entrainment factors for circadian clocks. BMC Mol Biol. 2006;7. doi:10.1186/1471-2199-7-5

69. Joseph NM, Mosher JT, Buchstaller J, Snider P, McKeever PE, Lim M, et al. The Loss of Nf1 Transiently Promotes Self-Renewal but Not Tumorigenesis by Neural Crest Stem Cells. Cancer Cell. 2008;13: 129–140. doi:10.1016/j.ccr.2008.01.003

70. Spyra M, Kluwe L, Hagel C, Nguyen R, Panse J, Kurtz A, et al. Cancer stem cell-like cells derived from malignant peripheral nerve sheath tumors. PLoS One. 2011;6. doi:10.1371/journal.pone.0021099

71. Livak KJ, Schmittgen TD. Analysis of relative gene expression data using real-time quantitative PCR and the 2-ΔΔCT method. Methods. 2001;25: 402–408. doi:10.1006/meth.2001.1262

72. Larionov A, Krause A, Miller WR. A standard curve based method for relative real time PCR data processing. BMC Bioinformatics. 2005;6: 62. doi:10.1186/1471-2105-6-62

73. Collins VP, Loeffler RK, Tivey H. Observations on growth rates of human tumors. Am J Roentgenol Radium Ther Nucl Med. 1956;76: 988–1000. Available: http://www.ncbi.nlm.nih.gov/pubmed/13362715

74. Gerlee P. The model muddle: In search of tumor growth laws. Cancer Research. 2013. pp. 2407–2411. doi:10.1158/0008-5472.CAN-12-4355

75. Benzekry S, Lamont C, Beheshti A, Tracz A, Ebos JML, Hlatky L, et al. Classical Mathematical Models for Description and Prediction of Experimental Tumor Growth. PLoS Comput Biol. 2014;10: e1003800. doi:10.1371/journal.pcbi.1003800

