## Supplemental Information for "“Temporal control of tumor growth in nocturnal mammals: impact of the circadian system”"

#### Supplemental Tables

**Suppl. Table 1:** qPCR Primer sequences.

**A.**

| **Transcript Name** | **Accession Number** | **Forward Primer (5´ → 3´)**<br>**Reverse Primer (5´ → 3´)** |
|---|---|---|
| *Tbp* | NM_013684.3 | AGAACAATCCAGACTAGCGCA [67]<br>GGGAACTTCACATCACAGCTC |
| *Bmal1* | NM_007489.3 | GCAGTGCCACTGACTACCAAGA [68]<br>TCCTGGACATTGCATTGCAT |
| *Per1* | NM_001159367.1 | CGGATTGTCTATATTTCGGAGCA<br>TGGGCAGTCGAGATGGTGT |
| *Pcyt-2* | NM_001347615.1 | GTCATCGCCGGCCTACACT<br>GAGTCCGCTCGTGCAGGTT |

**B.**

| **Transcript Name** | **Accession Number** | **TaqMan Gene Expression Assay (Applied Biosystems)** |
|---|---|---|
| *ChoKα* | NM_013490.3 | Mm00442760_m1 |
| *Lipin*1 | NM_001130412.1 | Mm00522212_m1 |

Summary of primers sequences for qPCR indicating transcript name, accession number and forward/reverse sequences.

**Suppl. Table 2: Primers sequences for genotyping of *Nf*1 and *Trp*53 genes.**

| **Genotyping of *Nf*1 and *Trp*53 genes** | | | |
|---|---|---|---|
| **Gene** | **Primer** | **Primer Sequence (5´ → 3´)** | **Bands pattern** |
| *Nf1* | Nf1 WT | TTCTGGCCTTATTGGACACC | **WT** (Nf1 COM- Nf1 WT): 374 bp<br>**MUT** (Nf1 COM- Nf1 MUT) 274 bp |
| | Nf1 COM | GCACAAAAGAGGCACTGGAT | |
| | Nf1 MUT | GGAGAGGTTTTTGCTTCCT | |
| *Trp53* | X7 | TATACTCAGAGCCGGCCT | **WT** (X7-X6.5): 450 bb<br>**MUT** (X7- NEO18) 550 bp |
| | X6.5 | ACAGCGTGGTGGTACCTTAT | |
| | NEO18 | CTATCAGGACATAGCGTTGG | |

Summary of primers sequences for genotyping of *Nf*1 and *Trp*53 genes indicating gene name, primer name, primer sequence and bands pattern. WT = wild type, MUT = mutant, bp = base pairs.

**Suppl. Table 3: Antibodies list.**

| Antibody | Host | Dilution | Catalogue |
|---|---|---|---|
| TUBULIN | Mouse | 1:1000 | Sigma (Cat# T9026) |
| SOX-2 | Rabbit | 1:200 | Chemicon (Cat# AB5603) |
| S-100 | Mouse | 1:250 | Sigma (Cat# S2532) |
| PER1 | Mouse | 1:100 | Abcam (Cat# ab3443) |
| GFAP | Rabbit | 1:250 | Dako (Cat# V5255) |
| VIMENTIN | Mouse | 1:750 | Sigma (Cat# V5255) |
| GLUTAMINE SYNTHASE | Mouse | 1:100 | Milipore (Cat# MAB302) |

Summary of antibodies used for immunochemistry indicating antibody name, host, dilution and catalogue number. GFAP = glial fibrillary acidic protein.

### SUPPLEMENTARY FIGURE 1

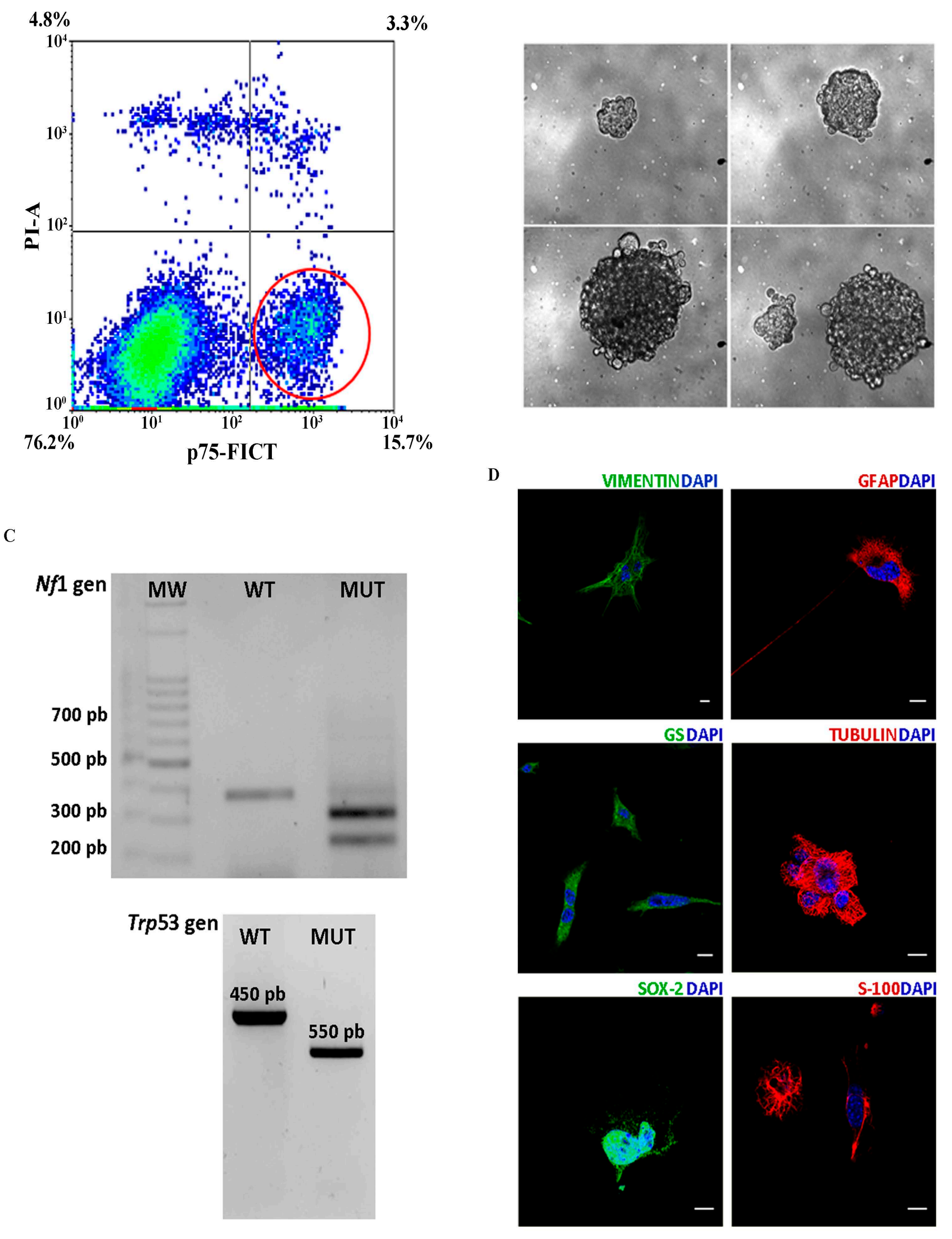

### SUPPLEMENTARY FIGURE 2

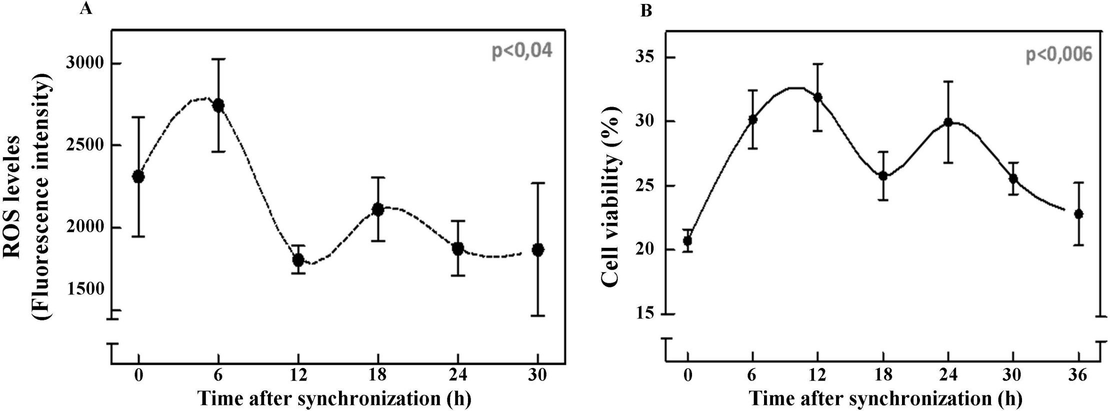

**SUPPLEMENTARY FIGURE 3**

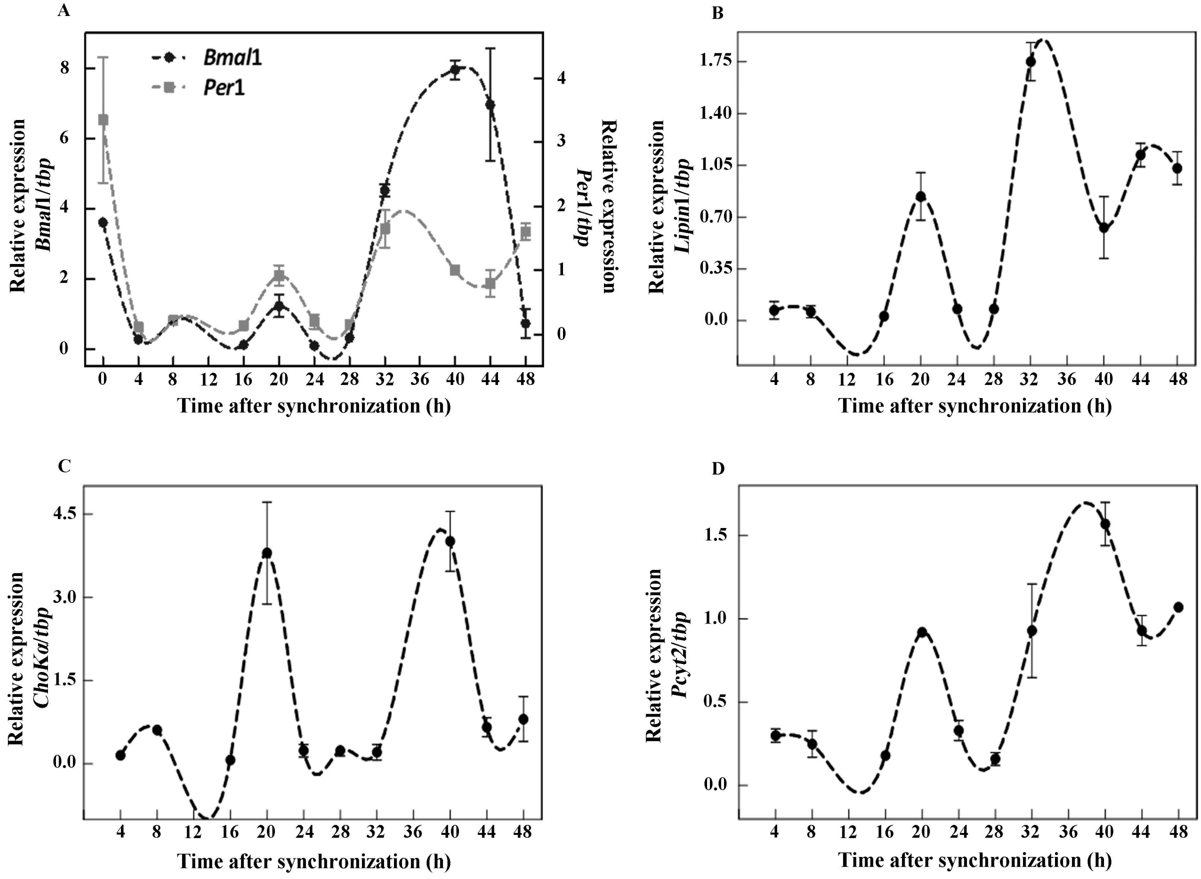

SUPPLEMENTARY FIGURE 4

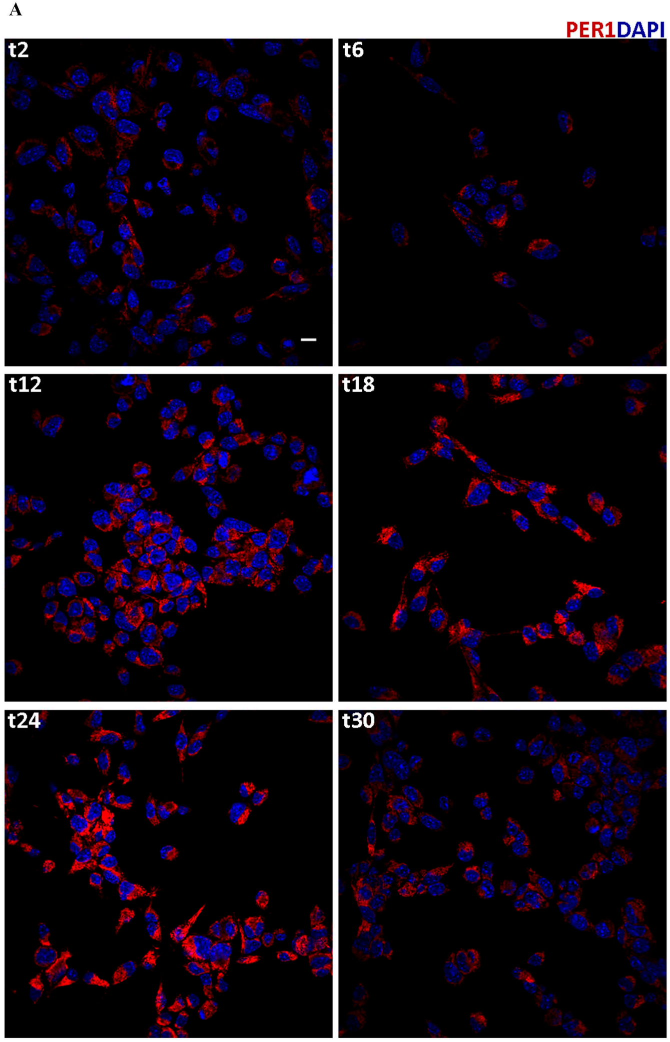


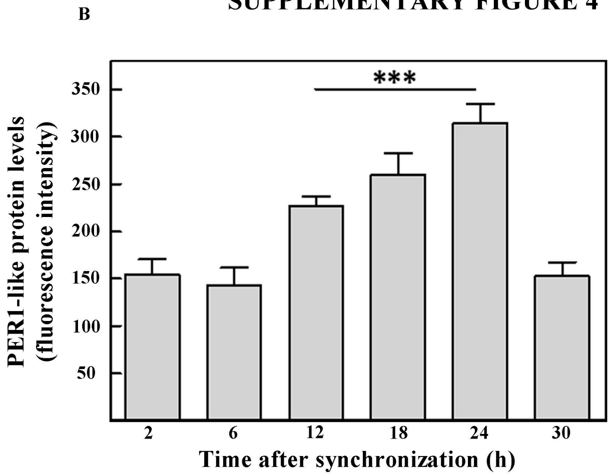

**SUPPLEMENTARY FIGURE 5**

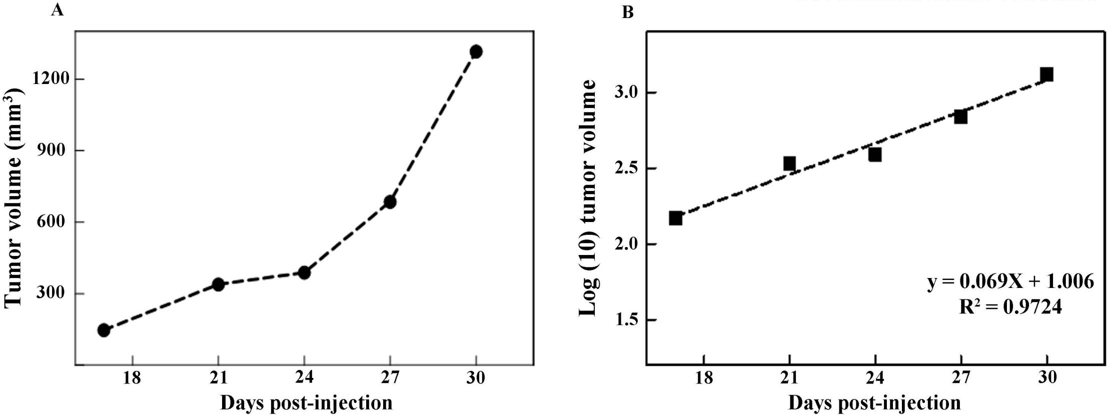

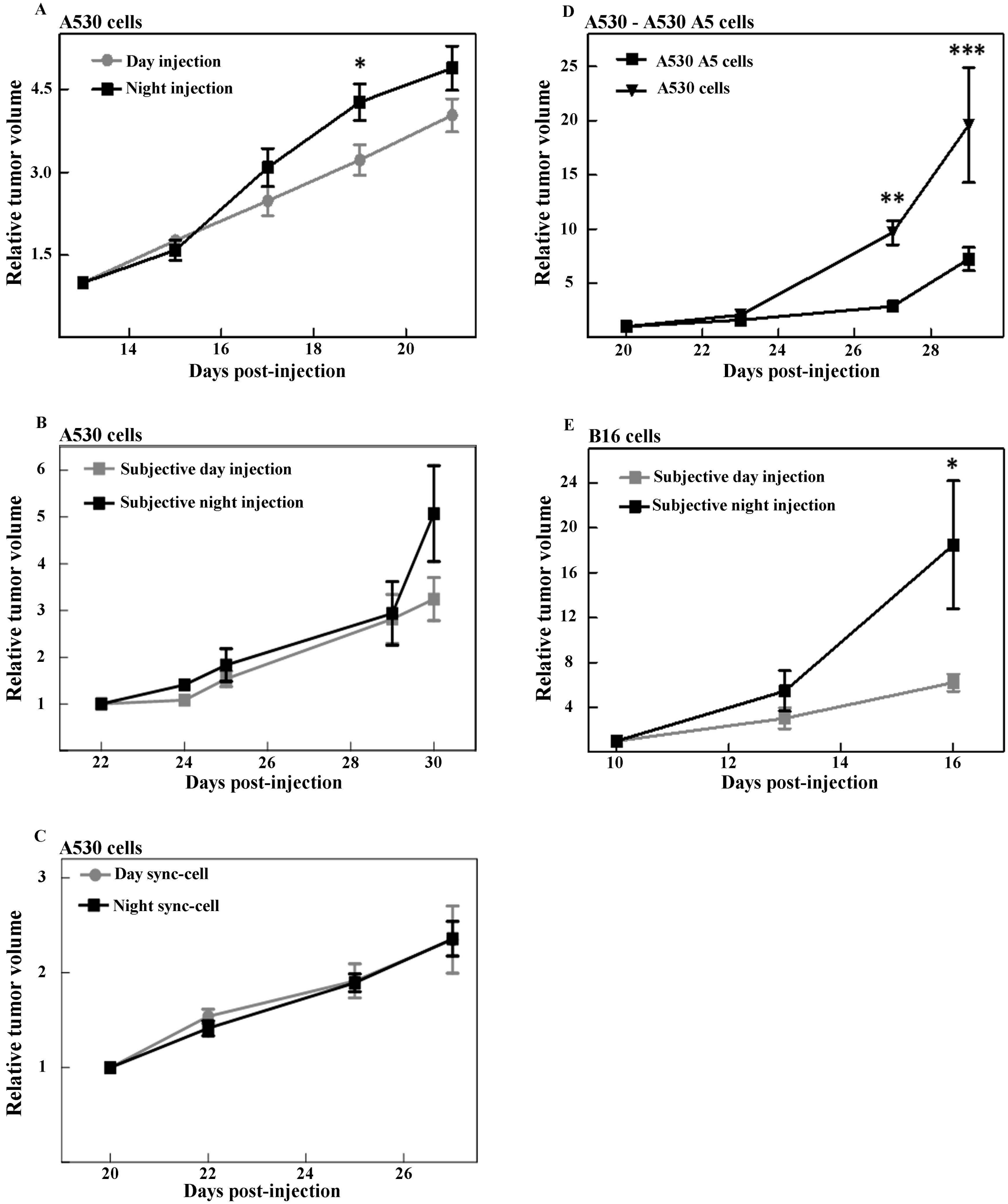
A
A530 cells
Day injection
Night injection
*
Relative tumor volume
Days post-injection
B
A530 cells
Subjective day injection
Subjective night injection
Relative tumor volume
Days post-injection
C
A530 cells
Day sync-cell
Night sync-cell
Relative tumor volume
Days post-injection
D
A530 - A530 A5 cells
A530 A5 cells
A530 cells
**
***
Relative tumor volume
Days post-injection
E
B16 cells
Subjective day injection
Subjective night injection
*
Relative tumor volume
Days post-injection

SUPPLEMENTARY FIGURE 7

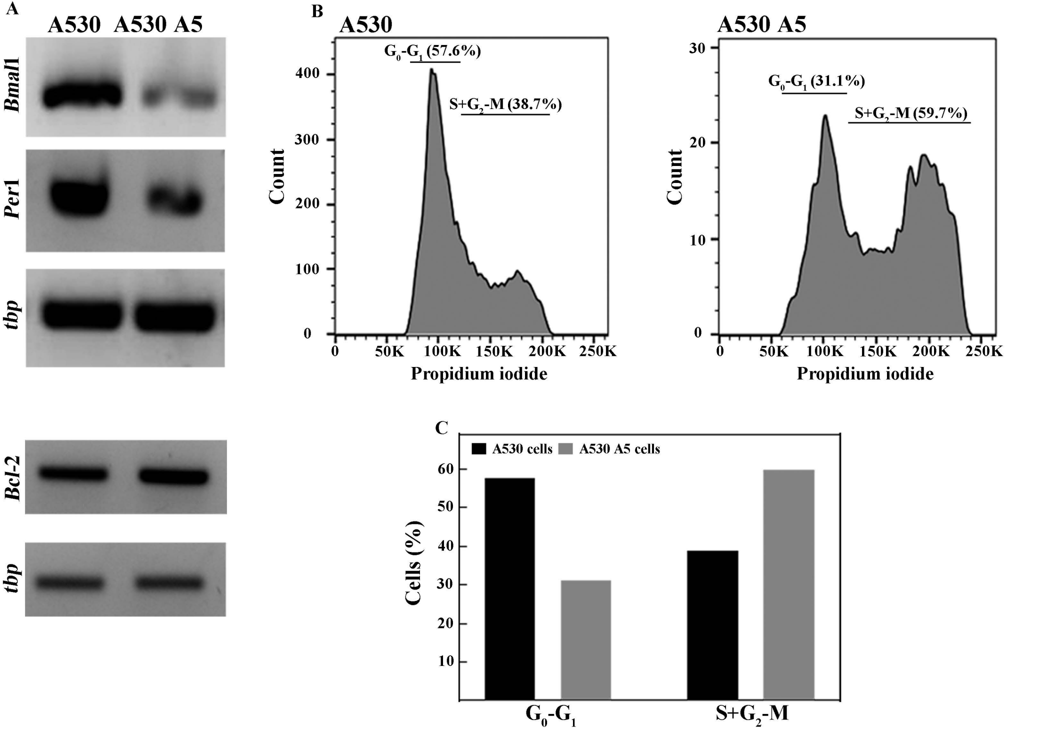

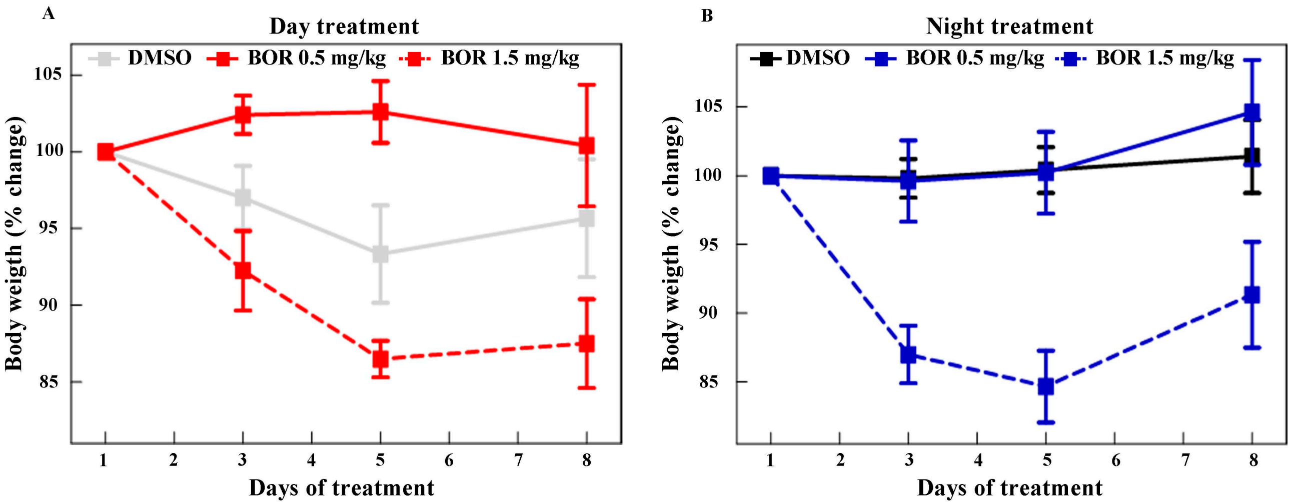
A
Day treatment
DMSO
BOR 0.5 mg/kg
BOR 1.5 mg/kg
105
100
95
90
85
Body weigth (% change)
1
2
3
4
5
6
7
8
Days of treatment
B
Night treatment
DMSO
BOR 0.5 mg/kg
BOR 1.5 mg/kg
105
100
95
90
85
Body weigth (% change)
1
2
3
4
5
6
7
8
Days of treatment

### Supplementary Figure Legends

**Suppl. Fig. 1: Characterization of A530 cell cultures A.** Cell sorting was used to isolate the p75$^{NGFR +}$ cells from a malignant peripheral nerve sheath tumor (MPNST). Representative flow cytometry dotplot showing that approximately 15.6% of cells express p75NGFR as shown in the lower panel on the right and marked with a red circle**. B.** Isolated p75$^{NGFR +}$ cells were grown in non-adherent conditions showing the capacity of spheroid forming. **C.** Genotyping of *Nf*1 and *Trp*53 genes of DNA obtained from A530 cells (MUT) and a control sample of C57BL/6 mouse tail (wild-type, WT) showing the loss of WT copy of both genes in A530 cells. **D**. Immunofluorescence in A530 cells with specific primary antibodies for the glial markers vimentin (upper panel, left column), glial fibrillary acidic protein (GFAP) (upper panel, right column) and glutamine synthetase (GS) (middle panel, left column), tubulin to denote the cellular cytoskeleton (middle panel, right column), and stem cell markers S-100 (lower panel, left column) and SOX-2 (lower panel, right column). Nuclei were stained with DAPI. Scale bar = 10 µm. MUT = mutant, WT = wild-type.

**Suppl. Fig. 2: Temporal variations in ROS levels (A) and susceptibility to Bortezomib chemotherapy (B) in A530 cell cultures. A.** Cells synchronized by culture medium exchange were collected at different times and the redox state of the cells was analyzed with the fluorescent probe 2′, 7′-Dichlorodihydrofluorescein diacetate to a final concentration of 2 µM as described in Methods. Cells were washed twice with PBS 1X and the fluorescence intensity was analyzed by flow cytometry. A significant temporal variation was observed in A530 cells ($p < 0.04$ by Kruskal-Wallis). **B**. Cells were synchronized by culture medium change and treated with Bortezomib (BOR, 500 nM) at different times post-synchronization for 36 h. A significant temporal variation was observed in levels of cell viability ($p < 0, 006$ by Kruskal-Wallis). The results are mean ± SEM of three independent experiments (n=5-6/group).

**Suppl. Fig. 3: Temporal variations in clock genes *Bmal*1 and *Per*1 (A), clock-controlled genes and/or lipid synthesizing enzyme genes *Lipin*1 (B), *CkoKα* (C) and *Pcyt*2 (D) mRNA expression in A530 cell cultures.** mRNA levels were assessed by RT-qPCR with RNA extracted from cells collected at different times during 48 h after synchronization by culture medium exchange at time 0; values were normalized according to the expression of the housekeeping gene *tbp*. Results showed a significant temporal variation in the mRNA expression for all different transcripts examined ($p < 0.05$ by Kruskal-Wallis).

**Suppl. Fig. 4: Immunocytochemistry of PER1-like protein in synchronized A530 cell cultures**. **A**. Cells were synchronized by culture medium exchange and collected at different times from 0 to 30 h after synchronization. Cultures were immunolabeled with specific primary antibody for PER1-like protein (red) and

DAPI (blue) for nuclear localization and visualized by confocal microscopy as described in Methods. Scale bar = 10 µm. **B**. Histograms indicating relative levels of PER1-like protein fluorescent intensity in A530 cells at different times post-synchronization. Data are mean ± SEM. Results revealed a significant temporal variation in levels of PER1-like protein by immunocytochemistry (***p < 0.0001 by Kruskal-Wallis), with highest levels at 12, 18 and 24 h after synchronization, differing from those at 2, 6 and 30 h.

**Suppl. Fig. 5: Estimation of tumor growth rate. A.** Synchronized A530 cells (1 x $10^5$ cells re-suspended in 10 µl of PBS) were injected into the sciatic nerve zone of C57BL/6 mice. Once tumors were palpable, their volumes (long, short and height dimension) were measured periodically with a caliper and plotted as a function of days post-injection. **B.** Linear regression of log-transformed tumor volumes as a function of days post-injection in which the slope of the straight line indicates the tumor growth rate.

**Suppl. Fig. 6: Relative Tumor Volume in C57BL/6 mice.** Relative tumor volume as a function of days post-injection of tumor cells under different conditions. **A**. A530 cells injected at the beginning of the day (gray line) or night (black line) in mice under a regular LD cycle. **B**. A530 cells injected at the beginning of the subjective day (gray line) or night (black line) in mice maintained in constant darkness (DD) for 72 h after synchronization by a regular LD cycle. **C**. A530 cell cultures synchronized at different times 12 h apart by culture medium exchange (day-sync cell: gray line; night-sync cell: black line) and injected in mice at the same time. **D**. A530 A5 cells (triangle) that had decreased *Bmal*1 expression by CRISPR/cas9 and their respective control cells (square) injected in mice. **E**. B16 melanoma cells injected at the beginning of the day (gray line) or night (black line) in mice under a regular LD cycle. Results are mean ± SEM of 2-3 experiments. The statistical analysis indicates: *p < 0.05, **p<.001 and ***p<.0001 by 2-way ANOVA with Bonferroni multiple comparison test. See Methods for estimation of the relative tumor volume.

**Suppl. Fig. 7: Characterization of A530 A5 cultures cells with *Bmal1* knocked-down. A.** RT-PCR for *Bmal*1, *Per*1 (a downstream target gene for *Bmal*1), *Bcl*-2 (gene codifying for an antiapoptotic protein) and the housekeeping gene *tbp* in A530 and A530 A5 cells. Results showed a significant decrease in *Bmal*1 and *Per*1 mRNAs and an increase in *Bcl*-2 mRNA in A530 A5 cells compared with A530 cells. **B.** Cells were collected 24 h after synchronization and the cell cycle was analyzed by flow cytometry with propidium iodide staining as described in Methods. Left panel shows data for A530 cells and right panel for A530 A5 cells denoting a higher level of S-$G_2$/M compared with control cells. **C.** Histograms indicating cell cycle distribution where A530 cells (black) present the highest percentage of cells arrested ($G_0$/$G_1$, 58%) whereas A530 A5 cells (gray) exhibited a higher percentage in S+$G_2$-M (59.7%) phases as compared with controls (38.7 %).

**Suppl. Fig. 8: Body weight (% change) of Bortezomib-treated mice.** Body weight of BOR-treated mice at the beginning of the day (A, red) or night (B, blue) were measured every day of treatment and reported as percentage change. Results show a decrease of ~15% in the body weight of animals treated with the high dose of BOR (1.5 mg/kg, dash lines) in both experimental conditions as compared with animals administered with the low dose of BOR (0.5 mg/kg, red and blue complete lines) or DMSO (vehicle, gray and black lines). BOR = Bortezomib.
